# LLM-powered automatic scoring tools for memory recall

**DOI:** 10.64898/2026.10.01.755994

**Authors:** Rayna S. Tang, Savannah J. Born, Angelique I. Delarazan, Phillip L. Demarest, Ata B. Karagoz, Zachariah M. Reagh

## Abstract

Free recall of naturalistic stimuli, such as films and stories, reveals what people remember and how they organize it. Scoring such recall requires matching each recalled utterance to the encoded stimulus, and doing this manually often makes large studies impractical. Existing automated methods mostly return similarity or aggregate scores rather than explicit matches between recalled and stimulus content. Here we introduce an open-source, model-agnostic pipeline that uses a large language model (LLM) to divide the stimulus and the recall into information units and to match units between the two texts. The output is a table of unit-to-unit matches with accuracy labels, from which standard analysis scripts compute recall measures. We validated the pipeline with four LLMs (Claude Sonnet 4.5, GPT-5.4, Gemini 2.5 Pro, and Gemini 2.5 Flash) on three datasets. On typed recall of four short stories, agreement between each model and two trained raters (φ = .73–.79) was close to the agreement between the raters themselves (φ = .77). A second run reproduced each model’s matches (φ = .91–.97) more closely than the raters agreed with each other. On spoken recall of 10 films, every model attributed recall to the correct film with high agreement. On uncorrected speech-to-text recall of a television episode, agreement varied more across models. The median cost was under $1 per participant for every model. These results suggest that the pipeline can serve as a reliable, low-cost alternative to manual scoring of free recall.

## Introduction

Episodic memory research increasingly uses naturalistic stimuli, such as films and spoken stories (Chen et al., 2017; Lee & Chen, 2022; Reagh & Ranganath, 2023). Unlike word lists or single images, these stimuli unfold continuously, as experience does in daily life (Nastase et al., 2020; Sonkusare et al., 2019). Memory for such stimuli is often tested with free recall, in which participants describe everything they remember in their own words. Because recall is unconstrained, participants’ recollections reveal what they retained and how they organized it. Recall content can be scored separately for gist and for peripheral details, which are forgotten at different rates (Sacripante et al., 2023). Recall order shows how memory is organized, for example in temporal order or by topic (Delarazan et al., 2026). What is recalled also depends on the structure of the material. Story structure shapes how much of a story is recalled and in what order (Mandler, 1978; Mandler & Johnson, 1977), and events with more causal or semantic connections to other events are recalled more often (Cohn-Sheehy et al., 2021; Lee & Chen, 2022). With these measures, researchers can compare memory across people and predict which information is most likely to be recalled.

However, to quantify narrative recall, researchers must score each recall against the encoded stimulus. Trained raters typically divide the stimulus and the recall into small scenes and then decide which stimulus scene each recall scene reflects (e.g., Delarazan et al., 2026; Lee & Chen, 2022; Sacripante et al., 2023). For memories of personal events, the Autobiographical Interview instead counts details within predefined categories, separating internal (episodic) details from external details (Levine et al., 2002). This kind of scoring is time-consuming, and large studies are often impractical as a result. Manual scoring is also open to rater bias, and in educational assessment, raters’ severity has been found to drift over long scoring periods (Congdon & MeQueen, 2000; Wilson & Hoskens, 2001). Similar drift may affect the scoring of recall. As datasets grow to hundreds of participants, manual scoring may limit which questions can be studied.

Several automated methods have been developed to reduce this burden. One approach compares the stimulus and the recall in a shared semantic space, built with topic models (Heusser et al., 2021) or sentence embeddings (Panela et al., 2025; Shen et al., 2023). These methods scale well, but they return a graded similarity for each event or topic. In other words, a similarity score does not show which stimulus detail a recalled statement corresponds to, or whether the statement is accurate. For autobiographical memory, fine-tuned language models score Autobiographical Interview details in close agreement with human raters (Klus et al., 2025; van Genugten & Schacter, 2024). These models count details within fixed categories that they learn from human-scored narratives, so a new scoring scheme would require new labeled data. Without additional training, large language models (LLMs) can segment narratives into events much as humans do (Michelmann et al., 2025; Panela et al., 2025). Their overall scores for story recall also correlate strongly with manual scores (Martinez, 2024). Moreover, an LLM has been used to judge which clauses of a story each participant recalled, and the recall probability of each clause closely matched that from manual scoring (Georgiou et al., 2025). However, these studies validated scores aggregated over a whole recall or over participants. It remains unclear whether an LLM-based tool can produce explicit, unit-by-unit matches between stimulus and recall, with accuracy labels, that agree with trained raters as closely as the raters agree with each other.

Here we introduce and validate an open-source, model-agnostic LLM pipeline for scoring free recall. In the segmentation step, the pipeline divides the encoded stimulus and the recall into information units of comparable size. These information units are minimal propositions that each contain one core assertion. In the matching step, the pipeline matches units between the two texts in either direction, labels each match as exact or loose, and flags incorrect information and meta-comments. The output is a table of unit-to-unit matches with explicit references to the original sentences. Accuracy, level of detail, temporal order, and errors can then be computed from this table with standard analysis scripts. The LLM only segments and matches. All scoring rules are applied afterward in code that researchers control. The prompt sets the matching criteria, so researchers can change them without new training data.

We tested the pipeline on three datasets with four LLMs from three providers (Claude Sonnet 4.5, GPT-5.4, Gemini 2.5 Pro, and Gemini 2.5 Flash). Dataset 1 tested the full pipeline on typed recall of four short expository stories (Born & Reagh, in prep.). Two trained raters matched the same segmented units as the models, so we compared model–human agreement with the agreement between the raters. Bayes factors tested whether model–human agreement was equivalent to human–human agreement. We also re-ran the matching step to test whether each model reproduced its own matches. Datasets 2 and 3 came with human scoring on predetermined units, so they tested the matching step alone. In Dataset 2, spoken recall of 10 short films (Lee & Chen, 2022) was matched against 628 sub-events from all films at once, and agreement was scored at the film level. In Dataset 3, speech-to-text recall of a television episode (Delarazan et al., 2023, 2026) was matched against 37 events and scored at the scene level. These transcripts retained filler words, false starts, and transcription errors.

In Dataset 1, model–human agreement was close to the agreement between the two raters, and Gemini 2.5 Pro did not fall below it in either matching direction. A second run reproduced each model’s matches more closely than the raters agreed with each other. In Dataset 2, all models attributed spoken recall to the correct film with high agreement. In Dataset 3, agreement varied more across models, and GPT-5.4 agreed most closely with the human scoring. The pipeline and analysis code are freely available at: https://github.com/raynatang/recall-scorer.

## Method

### The automated scoring pipeline

#### Overview

The pipeline is an open-source Python package built on LangChain, a framework that provides a common interface to LLMs from different providers. A single class (AutoScorer) provides both the segmentation step and the matching step (Figure 1A). The LLM only segments text into information units and matches these units, and all scores are computed afterward from the matches in analysis code. The Pydantic library checks each LLM response against a predefined JavaScript Object Notation (JSON) schema, and a response with missing fields, extra fields, or malformed JSON is rejected. All models received the same prompts and worked examples. We set the temperature to 0.1 for the Claude and Gemini models to reduce run-to-run variability. The GPT models ran at the provider’s default temperature, because LangChain’s OpenAI interface does not pass this setting to GPT-5 models. The full prompts and worked examples are given in the Supplementary Materials and in the code repository (https://github.com/raynatang/recall-scorer).

**Figure 1.**
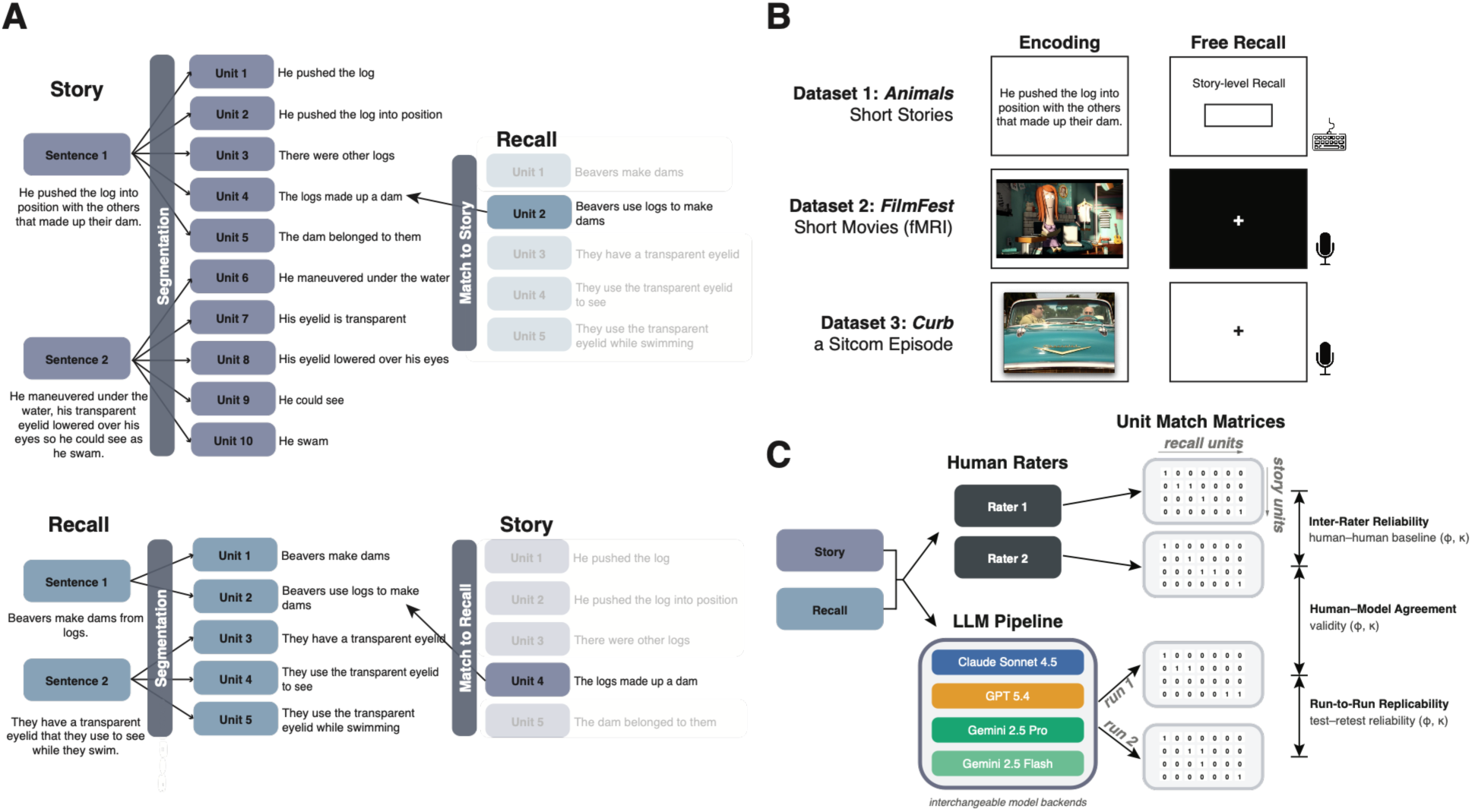
Overview of the pipeline and its validation. (A) The segmentation step divides each sentence of the story and of the recall into information units. In the matching step, each recall unit is matched to story units (story as template, top), or each story unit is matched to recall units (recall as template, bottom). (B) The three validation datasets, with the encoding materials and the recall format of each. (C) Evaluation design for Dataset 1. Two trained raters and each model matched the same story and recall units, and each model was run twice. Each scorer’s matches form a binary matrix of story units by recall units. Agreement between these matrices gives the human–human benchmark, human–model agreement, and run-to-run replicability, each measured with φ and κ.

#### Sentence tokenization

Each text is first split into sentences with the Punkt tokenizer in the Natural Language Toolkit (NLTK; Bird et al., 2009). Using a deterministic tokenizer instead of an LLM adds no cost and gives the same split on every run. The sentences keep their original order, and each sentence’s index is carried into all later outputs. Because Punkt relies on punctuation, long or unpunctuated passages, which are common in transcribed speech, remain single sentences. The segmentation prompt therefore tells the model to divide long sentences into clauses before it extracts information units.

#### Segmentation into information units

The segmentation step divides each sentence into information units, which are minimal factual statements that each contain one core assertion. Each unit has a simple structure of a subject, a verb, and an object or complement. The prompt’s rules turn each descriptive adjective into its own unit, so that “the big brown bats” yields “The bats are big” and “The bats are brown.” The prompt also asks the model to replace pronouns with their antecedents from the same sentence and to keep negations as negated units. Actions, locations, durations, goals, beneficiaries, and causal explanations each become separate units, as does each of several actions joined by a conjunction. Each unit must express a fact that the text states explicitly or directly implies, and the prompt forbids further inference, summaries, and interpretation. Every call includes the same seven worked examples (Supplementary Materials).

Sentences are sent to the model in batches of five, which keeps each prompt and response short. A response that fails the schema check stops the run with an error. The pipeline numbers the sentences and units of each valid response in order, so that C03_02, for example, is the second unit of the third sentence. For the Dataset 1 stories, the hand-annotated event-units were entered in place of sentences, so the first number identifies the event-unit. The output is a table with one row per unit, including its identifier, source sentence, and text.

#### Matching information units

The matching step finds the units of a recall and of the encoding text that express the same content. One unit list serves as the template, and for each template unit the model searches the other list, the pool, for units that match it. Matches are judged on meaning and context, not exact wording, so minor typos do not prevent a match. The recall-as-template direction shows what each recalled statement refers to and supports analyses of accuracy and errors. The story-as-template direction shows whether and where each encoded unit was recalled. For each matched template unit, the model returns its text and the text of each pool unit it matches, and it leaves out unmatched template units. The prompt favors a single exact match but allows several when pool units capture different facets of the template unit or jointly cover it.

Units are listed in the prompt without numbers, so the model reports each match by copying the unit’s text. Answering with unit numbers would make matching a multiple-choice task, and the ability of LLMs to link options to their labels varies greatly across models (Robinson et al., 2023). Moreover, LLMs favor particular option labels and positions independently of the options’ content (Pezeshkpour & Hruschka, 2023; Zheng et al., 2024). Numbering every unit would also change the prompt format, and LLM performance can be highly sensitive to such changes (Sclar et al., 2024). Copying text from the prompt, in contrast, is thought to rely on induction heads, which repeat token sequences that appeared earlier in the context (Feucht et al., 2025; Olsson et al., 2022).

The model labels each match as exact or loose. An exact match is accurate, with no meaningful distortion or missing detail. A loose match is true but vague, underspecified, or mildly distorted, and it may express a relationship that is implied rather than stated. A match that contains factually incorrect detail is flagged as wrong information, and a loose match cannot also be flagged as wrong information. With recall as template, the model also flags meta-comments, which describe the participant’s mental state (e.g., “I think I’m forgetting a part here”) or state inferences absent from the encoding text. A recall unit receives no match when no encoding unit supports it, including fabricated details, content from a different event, and pure meta-comments. When uncertain, the model is told to prefer no match over a loose match.

Long texts can produce responses that exceed a model’s output limit. Before matching, the pipeline therefore estimates the length of the response from the lengths of both unit lists. If the estimate exceeds the model’s output-token limit, the template units are split into batches, and each batch is matched against the full pool. In our datasets, batching was needed only in Dataset 2. If a response fails the schema check, its batch is split in half and resubmitted. A single template unit whose response still fails is dropped, and the failure is logged.

When the text of a matched pool unit occurs more than once in the pool, a follow-up LLM call identifies its source sentence. Given the template unit and the candidate sentences, the model chooses the sentence whose context best explains the match. A matched unit whose text still cannot be found in the pool is kept without an identifier and counted as unmatched (Supplementary Materials). The output is a single table with one row for each template unit, including unmatched ones, and an additional row for each further match. Each row holds the identifiers, source sentences, texts, and labels of both units.

#### Models

We tested whether the pipeline works with models from different providers, and we compared the models’ agreement with human scoring and their cost. For the main comparison, we chose one mid-tier model from each of three providers, in a similar price band. These were Claude Sonnet 4.5 (Anthropic; claude-sonnet-4-5-20250929), GPT-5.4 (OpenAI; gpt-5.4), and Gemini 2.5 Pro (Google; gemini-2.5-pro). Of these identifiers, only Claude’s names a dated model version. We accessed all models through the providers’ application programming interfaces (APIs) in June and July 2026. At July 2026 list prices, input and output tokens cost $3.00 and $15.00 per million for Claude Sonnet 4.5, $2.50 and $15.00 for GPT-5.4, and $1.25 and $10.00 for Gemini 2.5 Pro. We set each model’s output limit to the provider’s maximum, which was 64,000 tokens for Claude Sonnet 4.5, 128,000 for GPT-5.4, and 65,536 for the Gemini models. We did not test the providers’ premium models, because their higher prices would limit their use on large sets of recalls.

We also tested Gemini 2.5 Flash (Google; gemini-2.5-flash), a lower-cost model from the same family with list prices of $0.30 and $2.50 per million input and output tokens. We included it to test whether a model at a fraction of the price of the main models reaches similar agreement.

An earlier version of the main comparison used GPT-5.2 (gpt-5.2) in place of GPT-5.4. OpenAI announced the deprecation of GPT-5.2 during the study, so we replaced it with GPT-5.4 for all analyses in the main text. The replication and cost results for GPT-5.2 are reported in the Supplementary Materials.

### Validation datasets

We evaluated the pipeline on three datasets that each came with human scoring (Table 1; Figure 1B). Dataset 1 tested both steps of the pipeline on typed recall of written stories, and two trained raters provided the reference. Dataset 2 tested the matching step on spoken recall of 10 short films, which was matched against an existing annotation of all films at once. Dataset 3 tested the matching step on unedited speech-to-text transcripts of recall of a television episode, with one human rater’s scoring as the reference. The datasets also differed in unit length (Figure S1).

**Table 1.** Overview of the validation datasets.

|  | **Dataset 1: Animal stories** | **Dataset 2: FilmFest** | **Dataset 3: Curb** |
| --- | --- | --- | --- |
| Stimuli | 4 written expository stories (Born & Reagh, in prep.) | 10 short films (Lee & Chen, 2022; Lee et al., 2023) | 1 TV episode (Delarazan et al., 2026) |
| Encoding units | LLM-segmented (121–162 per story) | Human annotation (628 sub-events) | Human annotation (37 events) |
| Stimulus/annotation words | 588–602 per text (2,386 total) | 12,299 across 10 movies (572–2,151 per movie) | 1,419 across 37 events |
| Recall modality | Typed free recall | Spoken recall, manually transcribed | Spoken recall, speech-to-text |
| Recall units | LLM-segmented | Transcribed utterances (1–55 words) | Recall descriptions segmented during scoring |
| Recall words | M = 100 per story recall (18–257) | M = 4,836 per participant (2,280–9,131) | M = 1,543 per participant (254–5,885) |
| N (analyzed) | 33 | 15 | 84 |
| Reference standard | 2 trained raters | Human raters' film attribution | One human rater's event scoring |
| Pipeline stages | Segmentation + matching | Matching only | Matching only |
| Models | All 4 | All 4 | All 4 |

Encoding and recall units averaged about six words in Dataset 1 and about 20 words in Dataset 2. In Dataset 3, both varied widely, and recall descriptions were longer on average than event descriptions.

Table 1. word counts use the same tokenizer as the unit word counts (Figure S1), so the Dataset 1 text lengths differ slightly from the 583–594 words reported by Born and Reagh (in prep.). The Dataset 2 total excludes the two 151-word annotations of the filler cartoon shown before each scanning run. Recall words for Dataset 1 are given per story recall, and each participant recalled four stories. LLM = large language model.

#### Dataset 1: Animal stories

Dataset 1 comes from a study of how narrative and causal structure shape the recall of short texts (Born & Reagh, in prep.). The stimuli were four written expository texts about North American mammals, each about 600 words long (Table 1). Participants heard each text as a narrated video that showed the text one sentence at a time, and they then typed their recall of each text (procedure in the Supplementary Materials). The texts had been annotated by hand into event-units at the sentence or clause level (beaver, 67; bat, 48; fox, 54; opossum, 56), the level at which the original study scored recall. Of the 41 participants retained under the original study’s criteria, we excluded eight whose recalls included content from other stories, because the models matched each recall only against the units of its own story. Matching against several narratives at once was tested in Dataset 2. The analyzed sample therefore comprised 33 participants and 132 story recalls.

Gemini 2.5 Flash segmented the four texts in May 2025 and the recalls in June 2025, using an earlier version of the segmentation prompt. We kept these segmentations because the human raters had scored them, so the raters and all models matched the same units. The story units were checked by hand but not edited. The texts yielded 121 (bat), 136 (opossum), 151 (fox), and 162 (beaver) information units. Encoding units averaged 6.0 words (SD = 2.2, range 2–16), and recall units averaged 6.1 words (SD = 2.1, range 2–15). Typed recalls averaged 99.9 words per story (SD = 46.9, range 18–257). Two trained raters (Rater 1 and Rater 2) independently scored every recall (Born & Reagh, in prep.). For each recall unit, they entered the identifiers of the story units it reflected and flagged incorrect information and subjective statements (personal comments or unstated inferences). Each of the four models matched the same units in both directions, with recall as template and with story as template.

#### Dataset 2: FilmFest

Dataset 2 comes from the FilmFest study of memory for short films (Lee & Chen, 2022), and the data are available on OpenNeuro (ds004042; Lee et al., 2023). Fifteen participants watched 10 short films (M = 4.54 min, range 2.15–7.75 min) and then recalled them aloud in any order (procedure in the Supplementary Materials).

Human raters in the original study transcribed each recall by hand and divided it into utterances of no more than 50 words, based on pauses and changes of topic (Lee et al., 2023; Lee & Chen, 2022). The human raters classified each utterance by its content (e.g., recall of specific movie events or a general comment about a movie), and for utterances that recalled movie events, they identified the events described. The encoding units came from an existing annotation in which annotators divided each film into sub-events and described each one in their own words. We used one annotator’s annotation, which comprised 628 sub-event descriptions of the 10 films and the two cartoon presentations. The descriptions refer to characters with generic labels such as “the man” and “the woman,” which recur across films. The model received the descriptions without film labels, so the start of each film was marked only by the description of its title card.

Each participant’s utterances served as recall units without further segmentation (M = 22.5 words, SD = 12.4, range 1–55 by our word count), and spoken recalls averaged 4,836 words per participant (range 2,280–9,131). Each recall was matched, with recall as template, against the descriptions of all films at once, so the model had to tell films apart as well as sub-events within a film. For each utterance, we compared the film of the model’s matched sub-event with the film to which the human raters had attributed the utterance. The two cartoon presentations were coded as one non-film category, and utterances that the human raters did not link to a specific event, such as fillers and general comments, were treated as unmatched. Because character labels recurred across films and participants could recall the films in any order, this dataset tests whether the matching step can use narrative context rather than names or recall order. The text on both sides was clean, however, because the recalls were transcribed by hand and the annotation was written in full sentences.

#### Dataset 3: Curb

Dataset 3 comes from a study of how younger and older adults recall a television episode (Delarazan et al., 2026), which used the viewing task of Delarazan et al. (2023). Participants watched a 26-min episode of the sitcom *Curb Your Enthusiasm* (HBO; Season 1, Episode 7, “AAMCO”), in which several storylines are interleaved. They then recalled aloud everything they could remember about the episode, in any order. We scored the recalls of 84 participants (46 younger and 38 older adults). The recordings were transcribed with the Whisper speech-to-text model (Radford et al., 2022), and the transcripts were not proofread. They retain filler words, false starts, and recognition errors, and many consist of long stretches of text with little or no punctuation.

The encoding units were 37 event descriptions written by three annotators, who divided the episode at transitions between storylines and agreed on the final descriptions (Delarazan et al., 2026). Event descriptions averaged 38.4 words (SD = 36.8, range 3–200). As the human reference, one human rater segmented each transcript while scoring it. A segment was marked, without rewording, wherever the transcript recounted an annotated event or a detail unrelated to the episode, and each segment was labeled with the event it recounted, if any. These recall descriptions served as recall units without LLM segmentation. They averaged 53.7 words (SD = 58.6, range 2–585), and recalls averaged 1,543 words per participant (range 254–5,885). Because matches are reported as copied text, the model had to reproduce these long, often unpunctuated descriptions in its responses.

Each model matched each recall, with recall as template, against all 37 event descriptions. We compared the models’ event assignments with those of the human rater at the scene level, after grouping the 37 events into the 12 scenes defined in the annotation. Recall descriptions that the human rater did not assign to an event were treated as unmatched. This dataset therefore tests matching with a small set of candidates from one episode but with unedited speech-to-text input.

### Evaluation analyses

#### Segmentation consistency

Because no ground-truth segmentation exists, we tested whether the four models segmented the same text at a consistent level of detail. Each model segmented the typed recalls of Dataset 1 with the current prompt, and we counted the information units each model produced for each sentence that all four models segmented. For each model, we computed the Spearman correlation between the number of words and the number of units per sentence. For each pair of models, we computed the intraclass correlation (ICC) of units per sentence within each story recall with at least five sentences, using the one-way random-effects, single-measure form (ICC(1,1)), and we averaged it across recalls. We compared the number of units per sentence across models with a repeated-measures ANOVA (Greenhouse–Geisser corrected) and Holm-corrected paired t tests.

#### Match-matrix agreement

For each recall, we represented each scorer’s matches as a binary matrix with story units (or films or scenes) as rows and recall units as columns, in which 1 marks a link between the two (Figure 1C). Agreement between two scorers was the φ coefficient between their vectorized matrices. We tested each φ with a permutation test. One scorer’s matrix was shuffled 1,000 times, first by rows and then by columns, which preserves its sparsity, and φ was recomputed after each shuffle. The two-tailed p value was (k + 1)/(1,000 + 1), where k is the number of shuffled φ values at least as large in absolute value as the observed φ. Within each story recall (Dataset 1) or participant (Datasets 2 and 3), p values were corrected across all scorer pairs with the Benjamini–Hochberg false discovery rate (FDR) procedure, separately for each matching direction. In Dataset 1, we also repeated the analysis with event-units as rows and recall sentences as columns (Supplementary Materials). In Datasets 2 and 3, each match was credited to the film or scene that contained the matched encoding unit. We summarize each scorer pair by its mean φ and by the percentage of story recalls or participants with corrected p < .05.

#### Comparing model–human agreement with the human–human benchmark

To compare agreement between scorer pairs, we capped φ at ±.999, applied the Fisher z transformation, and averaged the z values within each participant. We compared scorer pairs with paired t tests and corrected the p values across all pairwise comparisons with the Benjamini–Hochberg procedure. Because the central question was whether model–human agreement was as high as human–human agreement, we also computed Bayes factors for the paired t tests (BF10; default Cauchy prior, r = .707). BF10 below 1/3 indicates moderate evidence for no difference, and BF10 above 3 indicates moderate evidence for a difference. In Datasets 2 and 3, which have a single human reference, we compared the four human–model pairs with one another.

#### Item-level agreement (Cohen’s κ)

As a measure closer to traditional recall scoring, we also computed Cohen’s κ on which story units each scorer judged to be recalled. For each story recall and scorer pair, we listed every story unit that at least one of the two scorers linked to the recall. Each scorer’s label for a unit was the unit’s identifier if that scorer linked it and “none” otherwise, and κ was computed over these paired labels. Units that neither scorer linked were not included. We computed κ for information units and for event-units, in which case linked units were collapsed into their event-unit. Values of κ were averaged within each participant and compared with the human–human benchmark with the same paired tests and Bayes factors as for φ. In Datasets 2 and 3, κ was computed over films and scenes, respectively. The κ results are reported in the Supplementary Materials.

#### Run-to-run replicability

Because LLM output can vary between runs, we tested whether the matching step reproduced its own output. For Dataset 1, we re-ran the matching step with recall as template for all five models, including GPT-5.2, about one month after the original run (June and July 2026). The inputs were identical, so differences between the runs reflect sampling variability or changes in the deployed models over that month. Because both runs used the same segmented recalls, they could be compared unit by unit. For each recall unit, we scored whether both runs linked it to the same story unit, counting units that neither run matched as agreements, and we also computed this rate over units that at least one run matched. We also scored whether both runs assigned the same match category (exact, loose, wrong information, meta-comment, or no match). Test–retest φ and κ between the two runs were computed as described above. Each model’s test–retest agreement was compared with the Rater 1–Rater 2 benchmark with the same Fisher z t tests and Bayes factors.

Analyses used Python 3.10.18 with pandas 2.3.1, NumPy 1.26.4, SciPy 1.15.3, statsmodels 0.14.5, and pingouin 0.6.0. The pipeline used LangChain 1.2.12 (langchain-anthropic 1.3.5, langchain-openai 1.1.11, and langchain-google-genai 4.2.1), Pydantic 2.12.5, and NLTK 3.9.1.

## Results

### Dataset 1: Animal stories

#### Segmentation granularity is consistent across models

We first tested whether the four models segmented recall sentences at a similar granularity. For every model, the number of information units per sentence increased with the number of words in the sentence (Spearman ρ = .75 for Claude Sonnet 4.5, .80 for GPT-5.4, .80 for Gemini 2.5 Pro, and .81 for Gemini 2.5 Flash; Figure 2A). However, across the 1,084 sentences that every model segmented, the models differed in how many units they produced (F(3, 3249) = 281.55, p < .001, Greenhouse–Geisser ε = .98, generalized η² = .011; Figure 2C). Claude Sonnet 4.5 produced the fewest units per sentence (M = 3.07, compared with 3.63–3.81 for the other models; Holm-corrected ps < .001, Cohen’s |d| = 0.21–0.28). The other three models also differed from one another, but these differences were small (GPT-5.4, M = 3.81; Gemini 2.5 Flash, M = 3.70; Gemini 2.5 Pro, M = 3.63; Holm-corrected ps ≤ .005, Cohen’s |d| ≤ 0.07). Pairs of models also agreed on the number of units in each sentence of a story recall (mean ICC = .68–.85 across the 120 recalls with at least five sentences; Figure 2B), and this agreement was significant for 73–96% of recalls. Agreement was lowest for pairs that included Claude Sonnet 4.5 (.68–.77) and highest among GPT-5.4 and the two Gemini models (.83–.85). Because this ICC counts a constant difference between two models as disagreement, Claude Sonnet 4.5’s coarser segmentation likely lowered its agreement with the other models. Thus, although Claude Sonnet 4.5 segmented more coarsely, all four models divided the same sentences in proportion to their length.

**Figure 2.**
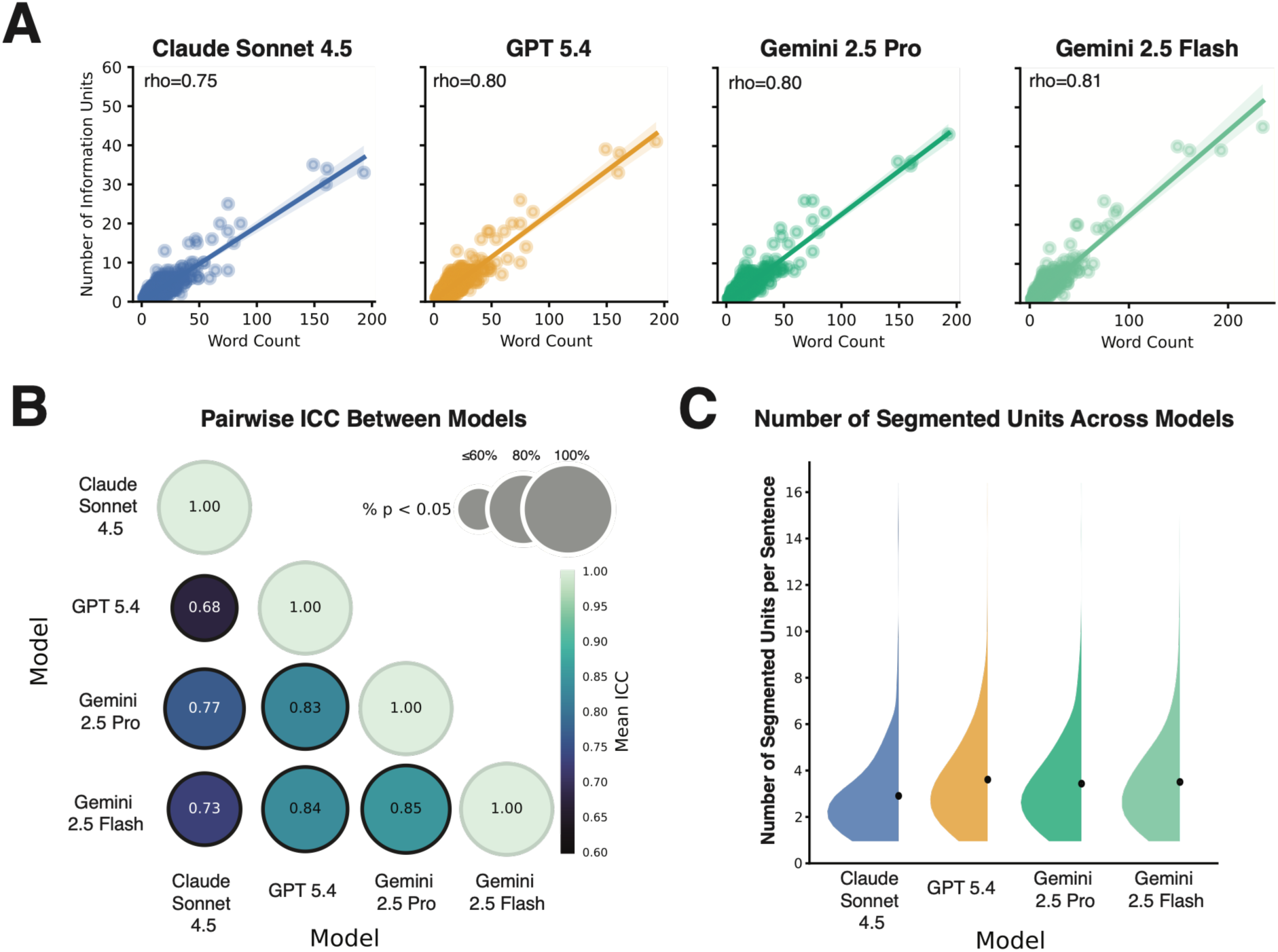
Segmentation of the Dataset 1 recalls by the four models. (A) Number of information units against the number of words for each recall sentence, with the linear fit and its 95% confidence band. Values of ρ are Spearman correlations. (B) Mean intraclass correlation (ICC(1,1)) of units per sentence between each pair of models, computed within each story recall with at least five sentences. Color shows the mean ICC, and circle size shows the percentage of recalls with p < .05. (C) Distribution of units per sentence across the 1,084 sentences that all four models segmented. Dots and bars show the mean ± SE.

#### Model–human agreement is close to the agreement between raters

We next compared each model’s matches with those of the two trained raters (Figure 3; Table S1). A match links one recall unit to one story unit. With recall as template, the raters made a mean of 20.3 (Rater 1) and 18.6 (Rater 2) matches per story recall, and the models made 17.8 (GPT-5.4) to 19.1 (Gemini 2.5 Pro and Gemini 2.5 Flash) matches (Claude Sonnet 4.5, 18.2; Figure 3A). The models thus made about as many matches as Rater 2 and slightly fewer than Rater 1. Agreement between the two raters averaged φ = .77 (SD = .12), and agreement between each rater and each model ranged from φ = .73 to .79 (Figure 3B). Every pair agreed significantly for all 132 story recalls (FDR-corrected permutation p < .05), except Rater 1 and Gemini 2.5 Flash, which agreed significantly for 131 recalls. The models agreed with one another more closely than with either rater (φ = .81–.87).

**Figure 3.**
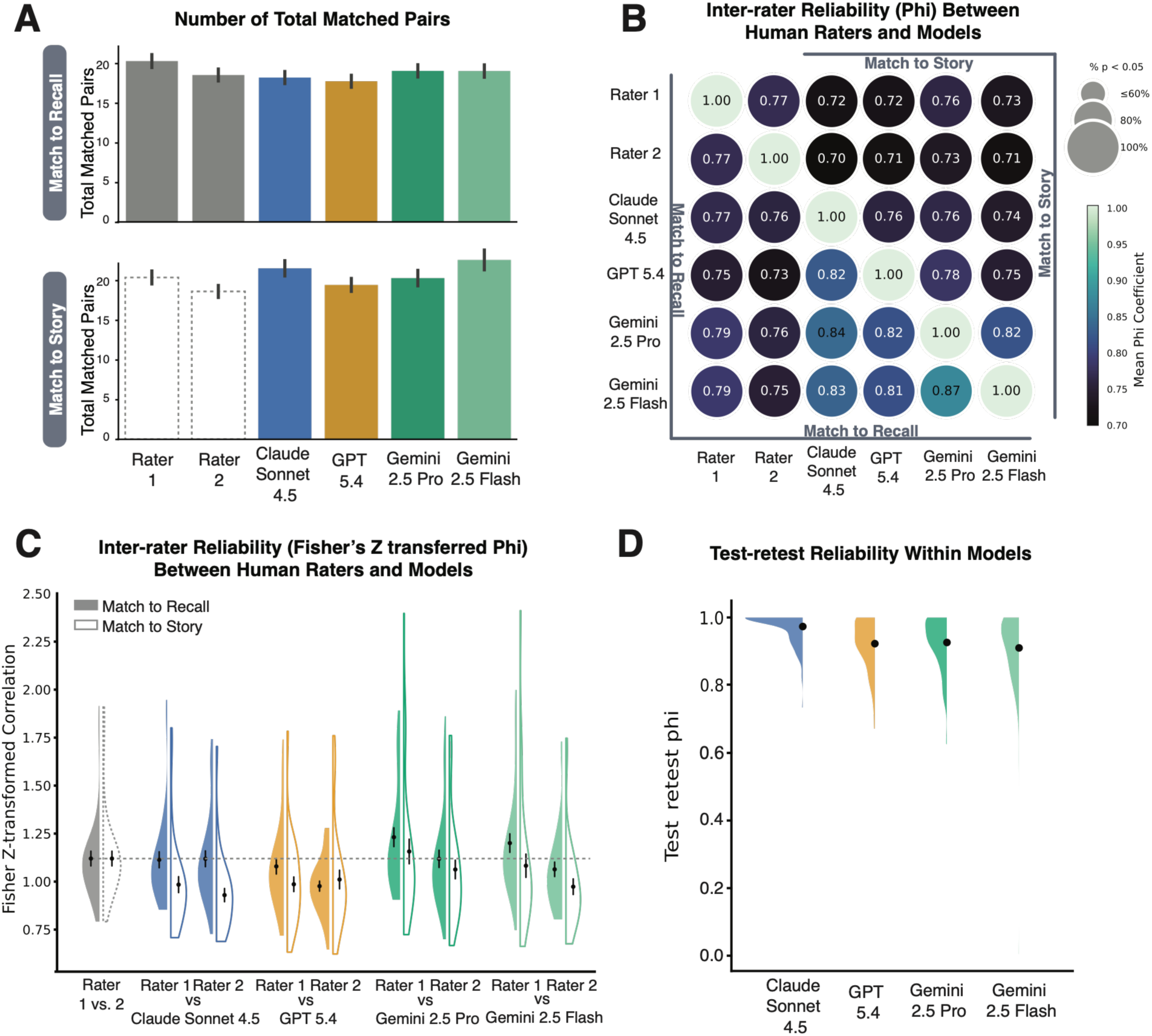
Agreement between the human raters and the models in Dataset 1. (A) Mean number of matches per story recall (N = 132) with recall as template (top, labeled “Match to Recall”) and with story as template (bottom, “Match to Story”). The raters scored only with recall as template, so their counts are repeated as dashed outlines in the bottom panel. (B) Mean φ between the match matrices of each pair of scorers, with recall as template below the diagonal and story as template above it. Color shows the mean φ, and circle size shows the percentage of story recalls with FDR-corrected p < .05. (C) Fisher z-transformed φ for each rater–model pair and for the two raters, averaged within participant (N = 33), with recall as template (filled) and story as template (outlined). The dashed line marks the mean agreement between the two raters. (D) Test–retest φ between the original and replication runs of each model, with recall as template (N = 132 story recalls). Bars in A and dots and bars in C and D show the mean ± SE.

Comparisons of Fisher z-transformed φ (Figure 3C) showed that only one human–model pair differed from the human–human benchmark after FDR correction. Agreement between Rater 2 and GPT-5.4 was lower than agreement between the two raters (t(32) = 3.90, corrected p = .004, BF10 = 64.1). Agreement between Rater 1 and Gemini 2.5 Pro was higher than the benchmark before correction but not after (t(32) = 2.32, uncorrected p = .027, corrected p = .069, BF10 = 1.91). The remaining six pairs did not differ from the benchmark (corrected ps ≥ .20, BF10 = 0.19–0.62). These Bayes factors gave moderate evidence for no difference (BF10 < 1/3) for Claude Sonnet 4.5 with both raters, GPT-5.4 with Rater 1, and Gemini 2.5 Pro with Rater 2, whereas they were inconclusive for Gemini 2.5 Flash with either rater (BF10 = 0.59 and 0.62).

With story as template, the models made 19.4 (GPT-5.4) to 22.5 (Gemini 2.5 Flash) matches per story recall (Claude Sonnet 4.5, 21.5; Gemini 2.5 Pro, 20.2; Figure 3A). Human–model agreement was lower in this direction (φ = .70–.76), as was agreement among the models (φ = .74–.82). Agreement fell below the human–human benchmark for Claude Sonnet 4.5 with both raters (corrected ps ≤ .009, BF10 ≥ 20.7) and for Gemini 2.5 Flash with Rater 2 (corrected p < .001, BF10 = 9.9 × 10³). GPT-5.4 also fell below the benchmark with both raters, although the Bayes factor for Rater 2 was weak (Rater 1, corrected p < .001, BF10 = 546; Rater 2, corrected p = .043, BF10 = 2.69). In contrast, Gemini 2.5 Pro did not differ from the benchmark with either rater (corrected ps ≥ .30, BF10 = 0.22 and 0.39), nor did Gemini 2.5 Flash with Rater 1 (corrected p = .60, BF10 = 0.23). We note that the raters scored recall only with recall as template. With story as template, the scorer instead searches each recall for up to 162 story units, so the lower agreement in this direction may partly reflect the different task rather than less accurate matching. Together, these results indicate that model–human agreement at the information-unit level was close to human–human agreement, especially when models and raters scored in the same direction. Gemini 2.5 Pro was the only model that did not fall below the benchmark in either direction. Agreement at the coarser event-unit level and item-level agreement (Cohen’s κ) are reported in the Supplementary Materials (Tables S2 and S3).

#### Model output replicates across runs more closely than raters agree

To test whether the matching step produces the same output across runs, we re-ran it on identical segmented inputs about one month after the original run (Figure 3D; Table S4). Of the 2,659 recall units, 90% (Gemini 2.5 Flash) to 96% (Claude Sonnet 4.5) were linked to the same story unit in both runs, counting units that neither run matched as agreements. The rates were similar when we counted only units that at least one run matched (90–96%). The two runs also assigned the same match category to 86% (Gemini 2.5 Pro and Gemini 2.5 Flash) to 95% (Claude Sonnet 4.5) of recall units. Test–retest agreement between the two runs’ match matrices was high for every model (φ = .91–.97). It was significant for every story recall, except one Gemini 2.5 Flash recall with a nearly empty match matrix (φ ≈ 0).

For every model, test–retest agreement exceeded the agreement between the two raters (φ = .77; ts(32) ≥ 10.38, corrected ps < .001, BF10 ≥ 9.8 × 10⁸). Claude Sonnet 4.5 was the most consistent model and exceeded each of the other three (ts(32) ≥ 6.81, corrected ps < .001, BF10 ≥ 1.4 × 10⁵). GPT-5.4, Gemini 2.5 Pro, and Gemini 2.5 Flash did not differ from one another (corrected ps ≥ .38, BF10 = 0.19–0.29). Test–retest κ showed the same pattern (Supplementary Materials, Table S5). Thus, a second run of the same model reproduced the first run more closely than a second trained rater reproduced the first rater’s scoring.

### Dataset 2: FilmFest

In Dataset 2, the human raters attributed each recall utterance to a film, so we evaluated the models’ matches at the film level. The human raters assigned a mean of 187.3 utterances per participant to a film, and the models matched 137.0 (Claude Sonnet 4.5) to 177.7 (Gemini 2.5 Pro) utterances to a sub-event (GPT-5.4, 175.8; Gemini 2.5 Flash, 166.2; Figure 4A; Table S6). Film-level agreement with the human raters was high for every model (φ = .84 for Claude Sonnet 4.5, .92 for GPT-5.4, .94 for Gemini 2.5 Pro, and .89 for Gemini 2.5 Flash; Figure 4B), and it was significant for all 15 participants. The models also agreed with one another (φ = .80–.92).

**Figure 4.**
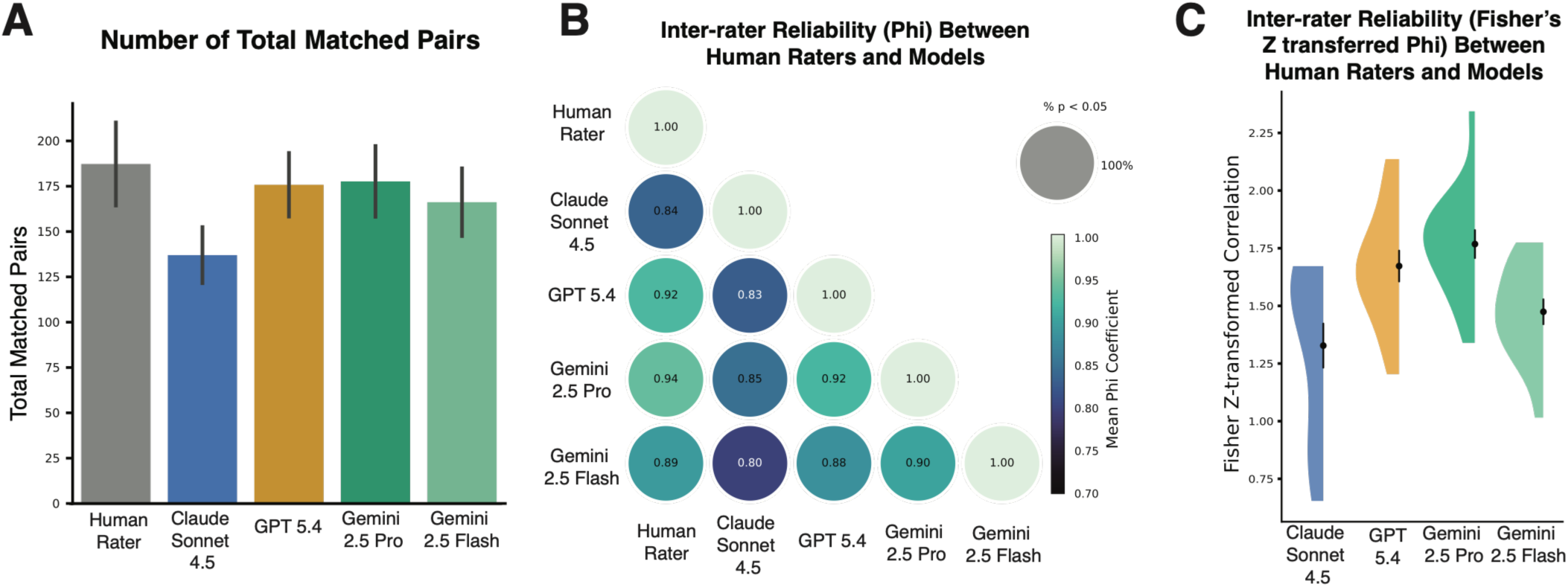
Film-level agreement in Dataset 2 (N = 15). (A) Mean number of recall utterances per participant that the human raters attributed to a film or that each model matched to a sub-event. (B) Mean φ between the film-by-utterance matrices of each pair of scorers. Color shows the mean φ, and circle size shows the percentage of participants with FDR-corrected p < .05. (C) Fisher z-transformed φ between each model and the human raters. Bars in A and dots and bars in C show the mean ± SE.

Pairwise comparisons of Fisher z-transformed φ (Figure 4C) showed that Gemini 2.5 Pro agreed with the human raters more closely than Claude Sonnet 4.5 (t(14) = 3.96, corrected p = .009, BF10 = 28.5) and Gemini 2.5 Flash (t(14) = 3.31, corrected p = .015, BF10 = 9.72). GPT-5.4 also agreed more closely than Claude Sonnet 4.5 (t(14) = 2.79, corrected p = .029, BF10 = 4.11). The other three comparisons (GPT-5.4 vs. Gemini 2.5 Pro, GPT-5.4 vs. Gemini 2.5 Flash, and Claude Sonnet 4.5 vs. Gemini 2.5 Flash) were not significant after correction (corrected ps ≥ .093), and their Bayes factors were inconclusive (BF10 = 0.56–1.31).

We note that the film-level criterion counts a match as correct whenever the chosen sub-event belongs to the correct film. It therefore does not detect errors among sub-events within a film, and these values may overstate agreement at the sub-event level. A human annotation of these recalls at the sub-event level would allow a stricter test. Film-level κ is reported in the Supplementary Materials (Table S7).

### Dataset 3: Curb

In Dataset 3, we evaluated agreement at the scene level, crediting each match to one of the 12 scenes into which the annotation groups the 37 events. The human rater matched a mean of 27.1 recall descriptions per participant to an event, whereas the models matched 15.5 (Claude Sonnet 4.5) to 24.6 (GPT-5.4) descriptions (Gemini 2.5 Pro, 21.6; Gemini 2.5 Flash, 17.8; Figure 5A; Table S8). Scene-level agreement with the human rater followed the same order (φ = .89 for GPT-5.4, .83 for Gemini 2.5 Pro, .75 for Gemini 2.5 Flash, and .70 for Claude Sonnet 4.5; Figure 5B). Agreement was significant for all 84 participants for GPT-5.4 and both Gemini models and for 82 participants for Claude Sonnet 4.5. Agreement among the models was lower than in Dataset 2 (φ = .61–.81).

**Figure 5.**
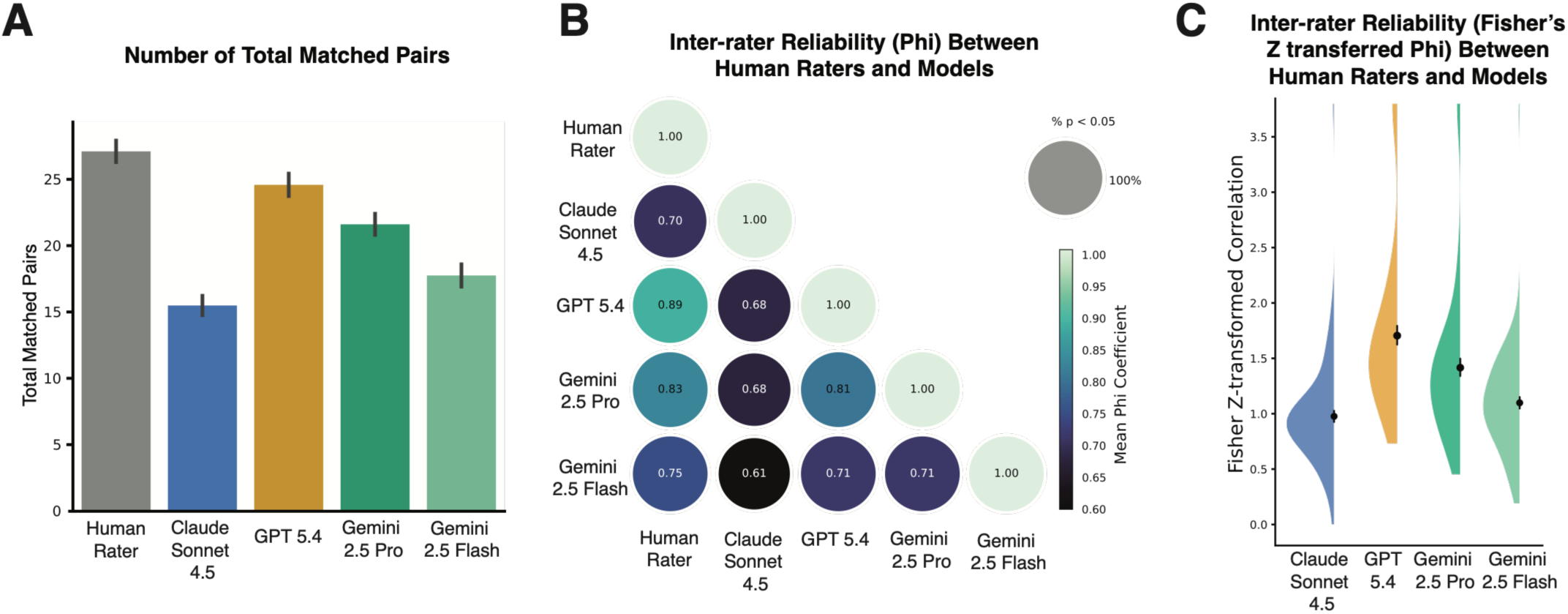
Scene-level agreement in Dataset 3 (N = 84). (A) Mean number of recall descriptions per participant that the human rater assigned to an event or that each model matched to an event. (B) Mean φ between the scene-by-description matrices of each pair of scorers. Color shows the mean φ, and circle size shows the percentage of participants with FDR-corrected p < .05. (C) Fisher z-transformed φ between each model and the human rater. Bars in A and dots and bars in C show the mean ± SE.

Pairwise comparisons of Fisher z-transformed φ (Figure 5C) showed that GPT-5.4 agreed with the human rater more closely than every other model. Its advantage was smallest over Gemini 2.5 Pro (t(83) = 3.15, corrected p = .003, BF10 = 11.3) and larger over Gemini 2.5 Flash and Claude Sonnet 4.5 (ts(83) ≥ 6.87, corrected ps < .001, BF10 ≥ 9.6 × 10⁶). Gemini 2.5 Pro in turn agreed more closely than Gemini 2.5 Flash (t(83) = 3.95, corrected p < .001, BF10 = 124) and Claude Sonnet 4.5 (t(83) = 4.95, corrected p < .001, BF10 = 4.0 × 10³). Claude Sonnet 4.5 and Gemini 2.5 Flash did not differ (t(83) = 1.78, corrected p = .078, BF10 = 0.55). Thus, on uncorrected speech-to-text recall, agreement varied more across models than in Datasets 1 and 2, and the models that matched fewer recall descriptions agreed less with the human rater.

The human reference also fixed the units that all scorers matched. These units varied widely in length (recall descriptions of 2–585 words and event descriptions of 3–200 words), whereas the matching prompt expects units of similar size, as the segmentation step produces in normal use. Agreement in Dataset 3 may therefore understate what the full pipeline achieves with segmented input. Scene-level κ is reported in the Supplementary Materials (Table S7).

## Discussion

The aim of the present study was to develop an open-source LLM pipeline that matches free recall to the encoded stimulus unit by unit, and to test its agreement with human scoring across three datasets. In Dataset 1, agreement between each model and each trained rater (φ = .73–.79) was close to the agreement between the two raters (φ = .77). With recall as template, only one of the eight model–rater pairs fell below this benchmark. Gemini 2.5 Pro did not fall below it in either matching direction. A second run of the matching step reproduced each model’s matches (φ = .91–.97) more closely than the two raters agreed with each other. In Dataset 2, every model attributed spoken recall of 10 films to the correct film with high agreement (φ = .84–.94). In Dataset 3, where uncorrected speech-to-text recall was matched to units of very uneven length, scene-level agreement was lower and varied more across models (φ = .70–.89). Taken together, these results suggest that the pipeline can serve as a reliable alternative to manual scoring of free recall.

Combining these results, our study shows that prompted LLMs can link recalled statements to specific units of the stimulus, and that these links can agree with trained raters about as well as the raters agree with each other. Prior automated methods compared recall with the stimulus using semantic embeddings (Heusser et al., 2021; Panela et al., 2025; Shen et al., 2023) or counted details within fixed categories (Klus et al., 2025; van Genugten & Schacter, 2024). Like these methods, the pipeline can score many recalls at low cost. However, its output is an explicit table of unit-to-unit matches with accuracy labels, and its matching criteria can be changed by editing the prompt. Earlier LLM studies of story recall validated aggregate scores, such as one overall score per recall (Martinez, 2024) or the recall probability of each story clause across participants (Georgiou et al., 2025). Here, agreement was assessed match by match within each recall, and in Dataset 1 it was compared with the agreement between two raters. Agreement also rose when matches were credited at the level of the hand-annotated event-units (human–model φ = .82–.87 vs. human–human φ = .86; Table S2). One reasonable interpretation is that scorers, whether human or model, disagreed mainly about which information unit within an event was recalled and less about which events were recalled.

The replication results also bear on the concern that raters may drift over long scoring periods. Each model reproduced its own matches about one month after the original run, which suggests that the matching step applies its criteria consistently over time. Similarly, in text annotation, the intercoder agreement of ChatGPT exceeded that of trained annotators (Gilardi et al., 2023).

However, consistency does not establish accuracy, because a model can repeat the same error. One might also argue that the models reached high agreement by matching only the most obvious units. However, with recall as template, the models made about as many matches per recall as Rater 2 (17.8–19.1 vs. 18.6), and only Rater 1 made more (20.3). Relatedly, agreement was lower with story as template (φ = .70–.76). Because the raters scored only with recall as template, the lower agreement in this direction may partly reflect a different task.

Our results also inform the choice of model. Gemini 2.5 Pro may be a good default. It was the only model that did not fall below the human–human benchmark in either matching direction in Dataset 1, and at the event-unit level its agreement with Rater 1 exceeded the benchmark (Table S2). It also had the highest film-level agreement in Dataset 2 (φ = .94), although it did not reliably differ from GPT-5.4 (BF10 = 0.59), and it ranked second in Dataset 3 (φ = .83). GPT-5.4 may be preferable for noisy speech-to-text recall. It agreed most closely with the human scoring in Dataset 3 (φ = .89) and exceeded Gemini 2.5 Pro (BF10 = 11.3). It was also the fastest model and had a lower median cost than Gemini 2.5 Pro. However, on typed recall it agreed less with Rater 2 than the two raters agreed with each other (BF10 = 64.1). Claude Sonnet 4.5 agreed with both raters at the level of the benchmark with recall as template, with moderate evidence for equivalence, and it was the most consistent model across runs. However, it segmented recall more coarsely than the other models and fell below the benchmark with story as template.

Within the Gemini family, Gemini 2.5 Pro agreed more closely with the human scoring than Gemini 2.5 Flash in Datasets 2 and 3 (BF10 = 9.72 and 124), whereas the two models were largely comparable in Dataset 1. Part of the deficit of Gemini 2.5 Flash reflects its own edits to the unit text it copied, which prevented those matches from being merged (Table S9). The ranking of models may also change with new releases, because performance on other text-analysis tasks has improved across successive model versions (Rathje et al., 2024). Because the pipeline is model-agnostic, new models can be benchmarked on the same datasets before they are adopted. Cost was low for every model. The median cost per participant was under $1 for every model and dataset, and the most expensive single participant cost $1.60 (Gemini 2.5 Pro, Dataset 2). Gemini 2.5 Pro was the slowest and most expensive model, whereas Gemini 2.5 Flash was the cheapest.

One limitation concerns how the pipeline records matches. The matching step identifies each match by copying the text of the matched unit, and the pipeline joins this copy back to the unit table. When a model altered the copied text, the match could not be merged and was scored as no match. This was rare in Dataset 1 (at most 0.17% of a model’s matches) but more common in Datasets 2 and 3, where units were longer, and it affected Gemini 2.5 Flash most (9.15% and 23.34%). Similar alterations, ranging from a changed word to truncated or combined text, have been reported when LLMs select quotes from qualitative research transcripts (Mehta et al., 2025). Because unmerged matches count as misses, they may understate the agreement of the affected models. A future version could re-submit batches that produce unmerged matches or accept near-exact copies through a more tolerant join. Adding an additional step at the beginning to clean up the input texts would also improve the stability of the output texts.

Another limitation is that all four models are closed commercial models. Their providers update and retire them, and OpenAI announced the deprecation of GPT-5.2 while this study was under way. Exact outputs may therefore not be reproducible later (Hussain et al., 2024; Palmer et al., 2024). Users of the pipeline should record the model version and date of each run and archive the raw outputs. Open-weight models that run locally would reduce this problem (Hussain et al., 2024), but we did not test them here. A third limitation is that the human references in Datasets 2 and 3 allowed only film-level and scene-level comparisons. These comparisons cannot detect errors within a film or scene, so agreement in Dataset 2 may overstate accuracy at the level of individual sub-events. In Dataset 3, the reference also fixed units of very uneven length (recall descriptions of 2–585 words), whereas the matching step is designed for units of similar size.

Agreement there may therefore understate what the full pipeline achieves when both texts are segmented first. [Unit-level human matching of these recalls is under way.]

The unit-level output of the pipeline makes several analyses feasible in large samples. For example, researchers could compare memory for gist and peripheral details, examine the temporal organization of recall, and track distortions in groups such as younger and older adults. With changes to the prompt, the same approach could also be adapted to other scoring schemes, such as the internal and external details of the Autobiographical Interview (Levine et al., 2002). In conclusion, an open-source LLM pipeline matched free recall to the encoded stimulus unit by unit and agreed with trained raters about as closely as the raters agreed with each other. Because it is inexpensive and consistent across runs, it may make unit-level scoring feasible in samples much larger than manual scoring allows.

## Supplementary Materials

### Unit lengths in the three datasets

**Figure S1.**
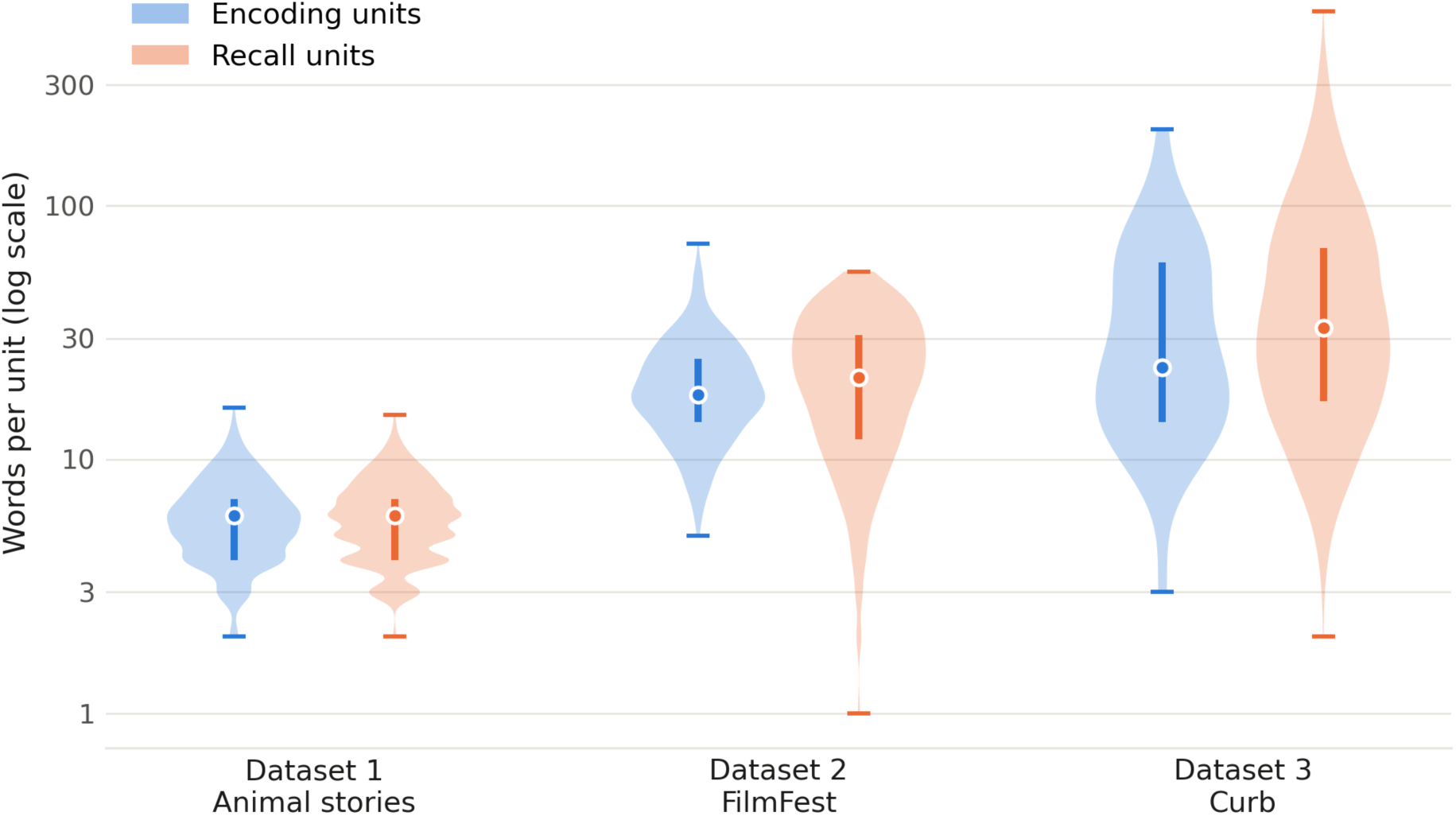
Word counts of the units matched in each dataset. Violins show the distribution of words per encoding unit (blue) and per recall unit (orange) on a log-scaled axis. Vertical bars mark the interquartile range, dots the median, and horizontal ticks the minimum and maximum. Dataset 1 includes the 33 analyzed participants. Unit lengths are similar within Datasets 1 and 2, whereas in Dataset 3 recall descriptions are longer and more variable than event descriptions.

### Datasets 1 and 2: Participants and procedure

The four texts of Dataset 1 crossed narrativity with causal structure, each at a high and a low level (Born, in prep.). The beaver text was high on both dimensions, the opossum text was low on both, the fox text was high only in narrativity, and the bat text was high only in causal structure. Drafts were generated with an LLM to keep style and level of detail similar across texts, and they were then fact-checked and revised to fit each condition. Forty-eight participants were recruited through the Washington University in St. Louis SONA pool or by word of mouth, and they received course credit or a $15 gift card for a session of about 50 min. Participants listened to AI-narrated videos of the four texts in random order (durations 3:59–4:38), which showed each text one sentence at a time, and they were told that they would later recall everything they remembered. After rating the stories and completing a 2-min arithmetic distractor task, participants typed their recall of each text, in random order and for at least 2 min per text. The response box showed only one line of text to discourage revision and to approximate spoken recall, and recall did not need to follow the order of the story. Under the original study’s criteria, six participants were excluded for low self-reported focus on the task and one for not following the recall instructions, which left 41 participants (M age = 20.1 years, range 18–27; 30 female).

In Dataset 2, 21 adults were recruited from the Princeton community, and six were excluded for excessive head motion during scanning, which left 15 participants (12 female; ages 20–33 years, M = 26.6; Lee & Chen, 2022). The 10 films comprised three animations and seven live-action films, and they were shown in two functional magnetic resonance imaging (fMRI) runs of five films each, with a 39-s filler cartoon before each run. Participants were encouraged to speak for at least 10 min during recall. The human raters classified each utterance as recall of specific movie events, a general comment about a movie, a memory search attempt, the end of recall, or speech unrelated to the task.

### Dataset 1: Agreement at the information-unit level

Table S1 lists the information-unit agreement shown in Figure 3B, with Bayes factors for the comparison with the human–human benchmark.

**Table S1.** Dataset 1: Mean phi coefficient (SD) between match matrices at the information-unit level (N *= 132 story recalls from 33 participants). BF10 compares each pair’s Fisher z-transformed φ with the Rater 1–Rater 2 benchmark (BF10 > 3, evidence for a difference; BF10 < 1/3, evidence for no difference). Every pair agreed significantly (FDR-corrected permutation p < .05) for all recalls, except Rater 1–Gemini 2.5 Flash with recall as template (131 of 132)*.

| Scorer pair | Recall as template, $\phi$ (SD) | BF10 | Story as template, $\phi$ (SD) | BF10 |
| --- | --- | --- | --- | --- |
| Rater 1–Rater 2 | .77 (.12) | — | .77 (.12) | — |
| Rater 1–Claude Sonnet 4.5 | .77 (.11) | 0.19 | .72 (.12) | 20.7 |
| Rater 2–Claude Sonnet 4.5 | .76 (.12) | 0.19 | .70 (.12) | $5.8 \times 10^4$ |
| Rater 1–GPT-5.4 | .75 (.13) | 0.25 | .72 (.13) | 546 |
| Rater 2–GPT-5.4 | .73 (.12) | 64.1 | .71 (.13) | 2.69 |
| Rater 1–Gemini 2.5 Pro | .79 (.11) | 1.91 | .76 (.12) | 0.22 |
| Rater 2–Gemini 2.5 Pro | .76 (.12) | 0.19 | .73 (.13) | 0.39 |
| Rater 1–Gemini 2.5 Flash | .79 (.12) | 0.62 | .73 (.14) | 0.23 |
| Rater 2–Gemini 2.5 Flash | .75 (.12) | 0.59 | .71 (.13) | $9.9 \times 10^3$ |
| Model–model (range) | .81–.87 | — | .74–.82 | — |

### Dataset 1: Agreement at the event-unit level

We repeated the Dataset 1 agreement analysis after collapsing story units into their hand-annotated event-units and recall units into their recall sentences (Table S2). Agreement rose for every pair (human–human φ = .86; human–model φ = .83–.87 with recall as template and .82–.87 with story as template), and it was significant for every recall. With recall as template, agreement between Rater 1 and Gemini 2.5 Pro exceeded the human–human benchmark (t(32) = 3.05, corrected p = .033, BF10 = 8.61), and no other pair differed from it (corrected ps ≥ .19, BF10 = 0.20–0.91). With story as template, no pair differed from the benchmark (corrected ps ≥ .12, BF10 = 0.20–1.76). Model–model agreement was .88–.93 with recall as template and .84–.89 with story as template. Scorers thus appear to disagree mainly about which information unit within an event-unit was recalled, rather than about which parts of the story were recalled.

**Table S2.**
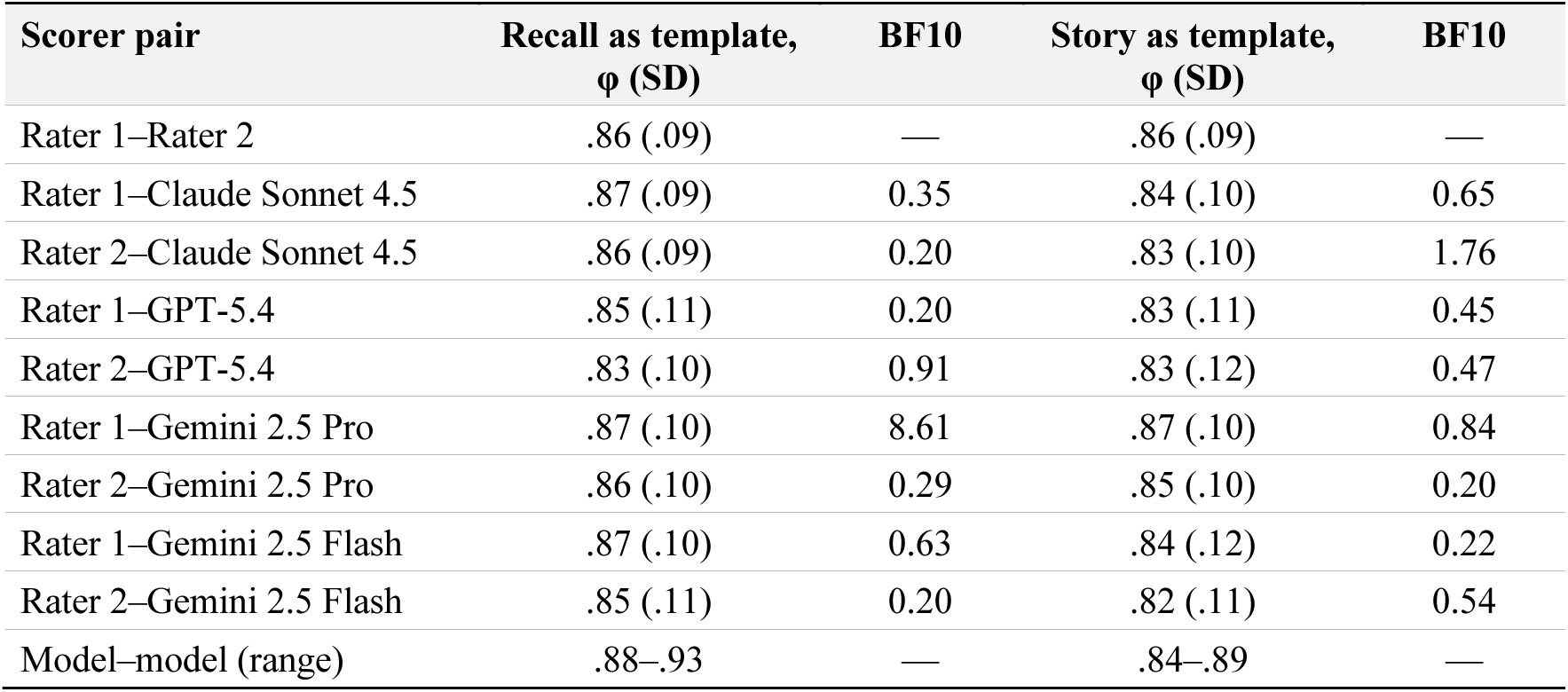
Dataset 1: Mean phi coefficient (SD) between match matrices at the event-unit level (N = 132 story recalls from 33 participants), with Bayes factors for the comparison with the Rater 1–Rater 2 benchmark (cf. Table S1). Every pair agreed significantly for all recalls.

### Dataset 1: Item-level agreement (Cohen’s κ)

Cohen’s κ on which story units each scorer judged recalled followed the φ results (Table S3). At the information-unit level, κ between the raters was .66, and κ between raters and models ranged from .59 to .67. GPT-5.4 agreed with both raters less than the raters agreed with each other (BF10 = 7.18 with Rater 1 and 4.6 × 10³ with Rater 2), and the evidence for Gemini 2.5 Flash with Rater 2 was weak (BF10 = 2.43). Bayes factors for the other five pairs ranged from 0.21 to 0.56. At the event-unit level, κ rose for every pair (human–human .77; human–model .72–.78), and only GPT-5.4 with Rater 2 still differed clearly from the benchmark (BF10 = 15.9).

**Table S3.** Dataset 1: Mean Cohen’s κ (SD) for recalled-unit judgments, with Bayes factors for the paired comparison with the Rater 1–Rater 2 benchmark (BF10 > 3, evidence for a difference; BF10 < 1/3, evidence for no difference). N = 33 participants.

| Scorer pair | Information-unit $\kappa$ (SD) | BF10 | Event-unit $\kappa$ (SD) | BF10 |
| --- | --- | --- | --- | --- |
| Rater 1–Rater 2 | .66 (.16) | — | .77 (.16) | — |
| Rater 1–Claude Sonnet 4.5 | .65 (.16) | 0.45 | .78 (.16) | 0.19 |
| Rater 2–Claude Sonnet 4.5 | .65 (.16) | 0.38 | .77 (.15) | 0.20 |
| Rater 1–GPT-5.4 | .62 (.17) | 7.18 | .74 (.17) | 1.59 |
| Rater 2–GPT-5.4 | .59 (.16) | $4.6 \times 10^3$ | .72 (.16) | 15.9 |
| Rater 1–Gemini 2.5 Pro | .67 (.16) | 0.26 | .78 (.16) | 0.22 |
| Rater 2–Gemini 2.5 Pro | .65 (.16) | 0.56 | .77 (.16) | 0.20 |
| Rater 1–Gemini 2.5 Flash | .67 (.17) | 0.21 | .78 (.16) | 0.23 |
| Rater 2–Gemini 2.5 Flash | .63 (.16) | 2.43 | .76 (.16) | 0.30 |

### Dataset 1: Run-to-run replication

Table S4 lists the replication results summarized in Figure 3D.

**Table S4.** Dataset 1: Run-to-run replication of the matching step (original vs. replication run; N = 132 story recalls from 33 participants; 2,659 recall units per model). Same match = proportion of recall units linked to the same story unit in both runs (units unmatched in both runs count as agreement). Same category = proportion assigned the same match category. Test–retest φ = mean phi (SD) between the two runs’ match matrices. Sig. = % of recalls with FDR-corrected permutation p < .05. BF10 compares Fisher z-transformed test–retest φ with the Rater 1–Rater 2 benchmark.

| Scorer pair | Same match | Same category | Test–retest $\phi$ (SD) | Sig. (%) | BF10 vs. benchmark |
| --- | --- | --- | --- | --- | --- |
| Rater 1–Rater 2 (benchmark) | — | — | .77 (.12) | 100 | — |
| Claude Sonnet 4.5 | .96 | .95 | .97 (.05) | 100 | $6.8 \times 10^{15}$ |
| GPT-5.4 | .91 | .89 | .92 (.07) | 100 | $1.6 \times 10^{10}$ |
| Gemini 2.5 Pro | .92 | .86 | .93 (.08) | 100 | $2.2 \times 10^{10}$ |
| Gemini 2.5 Flash | .90 | .86 | .91 (.11) | 99 | $9.8 \times 10^8$ |

### Dataset 1 replication: Test–retest κ

Test–retest κ between the original and replication runs was .96 (Claude Sonnet 4.5), .87 (GPT-5.4), .88 (Gemini 2.5 Pro), and .86 (Gemini 2.5 Flash) at the information-unit level, compared with .66 between the two raters (Table S5). At the event-unit level, the corresponding values were .97, .91, .92, and .92, compared with .77 between the raters. Every model exceeded the human–human benchmark at both levels (information-unit level, ts(32) ≥ 12.98, BF10 ≥ 2.6 × 10¹¹; event-unit level, ts(32) ≥ 8.95, BF10 ≥ 3.3 × 10⁷).

**Table S5.** Dataset 1: Test–retest Cohen’s κ (SD) between the original and replication runs, with Bayes factors for the paired comparison with the Rater 1–Rater 2 benchmark. N = 132 story recalls from 33 participants.

| Scorer pair | Information-unit $\kappa$ (SD) | BF10 | Event-unit $\kappa$ (SD) | BF10 |
| --- | --- | --- | --- | --- |
| Rater 1–Rater 2 (benchmark) | .66 (.16) | — | .77 (.16) | — |
| Claude Sonnet 4.5 (test–retest) | .96 (.08) | $5.3 \times 10^{17}$ | .97 (.06) | $2.6 \times 10^{12}$ |
| GPT-5.4 (test–retest) | .87 (.13) | $2.6 \times 10^{11}$ | .91 (.11) | $3.3 \times 10^7$ |
| Gemini 2.5 Pro (test–retest) | .88 (.12) | $7.9 \times 10^{12}$ | .92 (.11) | $4.0 \times 10^7$ |
| Gemini 2.5 Flash (test–retest) | .86 (.16) | $4.2 \times 10^{12}$ | .92 (.11) | $1.0 \times 10^8$ |

### Dataset 2: Film-level agreement

Table S6 lists the film-level results shown in Figure 4.

**Table S6.** Dataset 2: Film-level agreement between each model and the human raters (N = 15). Matches *= mean number of recall utterances per participant assigned to a film (human raters) or matched to a sub-event (models). φ = mean phi (SD) between the model’s and the human raters’ film-by-utterance matrices. Sig. = % of participants with FDR-corrected permutation p < .05*.

| Scorer | Matches | $\phi$ with human (SD) | Sig. (%) |
| --- | --- | --- | --- |
| Human raters | 187.3 | — | — |
| Claude Sonnet 4.5 | 137.0 | .84 (.12) | 100 |
| GPT-5.4 | 175.8 | .92 (.04) | 100 |
| Gemini 2.5 Pro | 177.7 | .94 (.03) | 100 |
| Gemini 2.5 Flash | 166.2 | .89 (.05) | 100 |

### Datasets 2 and 3: Item-level agreement (Cohen’s κ)

In Dataset 2, film-level κ was near ceiling for every model (.98–.99), and the models did not differ (corrected ps ≥ .87, BF10 = 0.26–0.40; Table S7). In Dataset 3, scene-level κ followed the φ results (GPT-5.4, .93; Gemini 2.5 Pro, .85; Gemini 2.5 Flash, .76; Claude Sonnet 4.5, .70).

GPT-5.4 exceeded Gemini 2.5 Pro (t(83) = 3.96, corrected p < .001, BF10 = 128), Gemini 2.5 Flash, and Claude Sonnet 4.5 (ts(83) ≥ 6.85, corrected ps < .001, BF10 ≥ 8.9 × 10⁶). Gemini 2.5 Pro exceeded Gemini 2.5 Flash (t(83) = 2.70, corrected p = .010, BF10 = 3.55) and Claude Sonnet 4.5 (t(83) = 5.67, corrected p < .001, BF10 = 6.7 × 10⁴). Claude Sonnet 4.5 and Gemini 2.5 Flash did not differ (t(83) = 1.83, corrected p = .071, BF10 = 0.59).

**Table S7.**
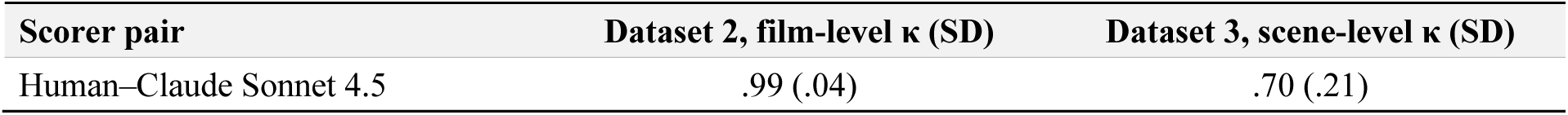
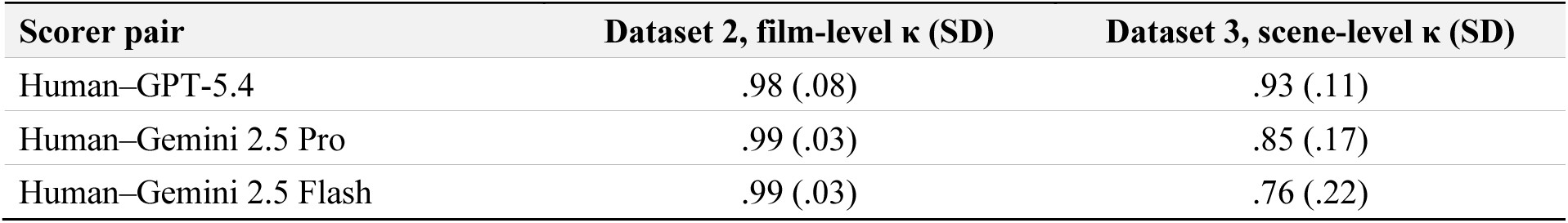
Datasets 2 and 3: Mean Cohen’s κ (SD) between each model and the human reference, computed over films (Dataset 2; N = 15) and scenes (Dataset 3; N = 84).

### Dataset 3: Scene-level agreement

Table S8 lists the scene-level results shown in Figure 5.

**Table S8.** Dataset 3: Scene-level agreement between each model and the human rater’s scoring (N = 84). Matches = mean number of recall descriptions per participant matched to an event. φ = mean phi (SD) between the model’s and the human rater’s scene-by-description matrices. Sig. = % of participants with FDR-corrected permutation p < .05.

| Scorer | Matches | $\phi$ with human (SD) | Sig. (%) |
| --- | --- | --- | --- |
| Human rater | 27.1 | — | — |
| Claude Sonnet 4.5 | 15.5 | .70 (.17) | 98 |
| GPT-5.4 | 24.6 | .89 (.09) | 100 |
| Gemini 2.5 Pro | 21.6 | .83 (.13) | 100 |
| Gemini 2.5 Flash | 17.8 | .75 (.17) | 100 |

### Matches that could not be merged back

The matching step identifies each match by copying the text of the matched unit, and the pipeline then joins this text back to the unit table. When a model altered the copied text, the join failed, and the match was kept in the output but scored as no match in every analysis. Such matches were rare in Dataset 1 (at most 0.17% of a model’s matches) and more common in Datasets 2 and 3 (Table S9). Gemini 2.5 Flash was affected most (9.15% of its matches in Dataset 2 and 23.34% in Dataset 3), followed by Gemini 2.5 Pro in Dataset 3 (4.42%) and Claude Sonnet 4.5 in Dataset 2 (3.16%).

**Table S9.** Matches whose copied text could not be joined back to the unit table, as a percentage of each model’s matches, with the maximum number for a single participant in parentheses. Dataset 1 uses the original run and the 33 analyzed participants.

| Dataset | Claude Sonnet 4.5 | GPT-5.4 | Gemini 2.5 Pro | Gemini 2.5 Flash |
| --- | --- | --- | --- | --- |
| Dataset 1 (animal stories) | 0.08 (2) | 0.17 (1) | 0.00 (0) | 0.12 (2) |
| Dataset 2 (FilmFest) | 3.16 (14) | 1.82 (9) | 0.82 (6) | 9.15 (40) |
| Dataset 3 (Curb) | 1.14 (2) | 0.24 (2) | 4.42 (6) | 23.34 (24) |

### Computational cost and runtime

We logged token usage and wall-clock time for every model call. For each participant, we summed input tokens, output tokens, and processing time across successful calls and estimated cost from each model’s list price per million input/output tokens (Claude Sonnet 4.5, $3.00/$15.00; GPT-5.4, $2.50/$15.00; Gemini 2.5 Pro, $1.25/$10.00; Gemini 2.5 Flash, $0.30/$2.50; GPT-5.2, $1.75/$14.00). These estimates ignore prompt-caching discounts and exclude failed calls, which we report separately below. For Dataset 1, we used the replication run, whose log covers every model on the same 43 participants.

The median cost per participant was below one US dollar for every model in every dataset (Figure S2). The highest cost for a single participant was $1.60 (Gemini 2.5 Pro in Dataset 2). GPT-5.4 was the fastest model in every dataset (median 27 s per participant in Datasets 1 and 3 and 176 s in Dataset 2), and Gemini 2.5 Pro was the slowest (121 s, 475 s, and 151 s in Datasets 1, 2, and 3). Gemini 2.5 Pro was also the most expensive model (median $0.22, $0.80, and $0.18 per participant), whereas Gemini 2.5 Flash was the cheapest ($0.07, $0.25, and $0.06). GPT-5.4 cost less than Claude Sonnet 4.5 in every dataset ($0.10 vs. $0.11, $0.55 vs. $0.61, and $0.07 vs. $0.09). Both Gemini models produced several times more output tokens than Claude Sonnet 4.5 and GPT-5.4. In Dataset 1, for example, the median output was 20,322 tokens for Gemini 2.5 Pro and 25,356 for Gemini 2.5 Flash, compared with 5,198 for Claude Sonnet 4.5 and 4,792 for GPT-5.4. Reasoning tokens made up 64–78% of the Gemini models’ output across datasets. Gemini 2.5 Flash was nonetheless the cheapest model because of its low list price.

**Figure S2.**
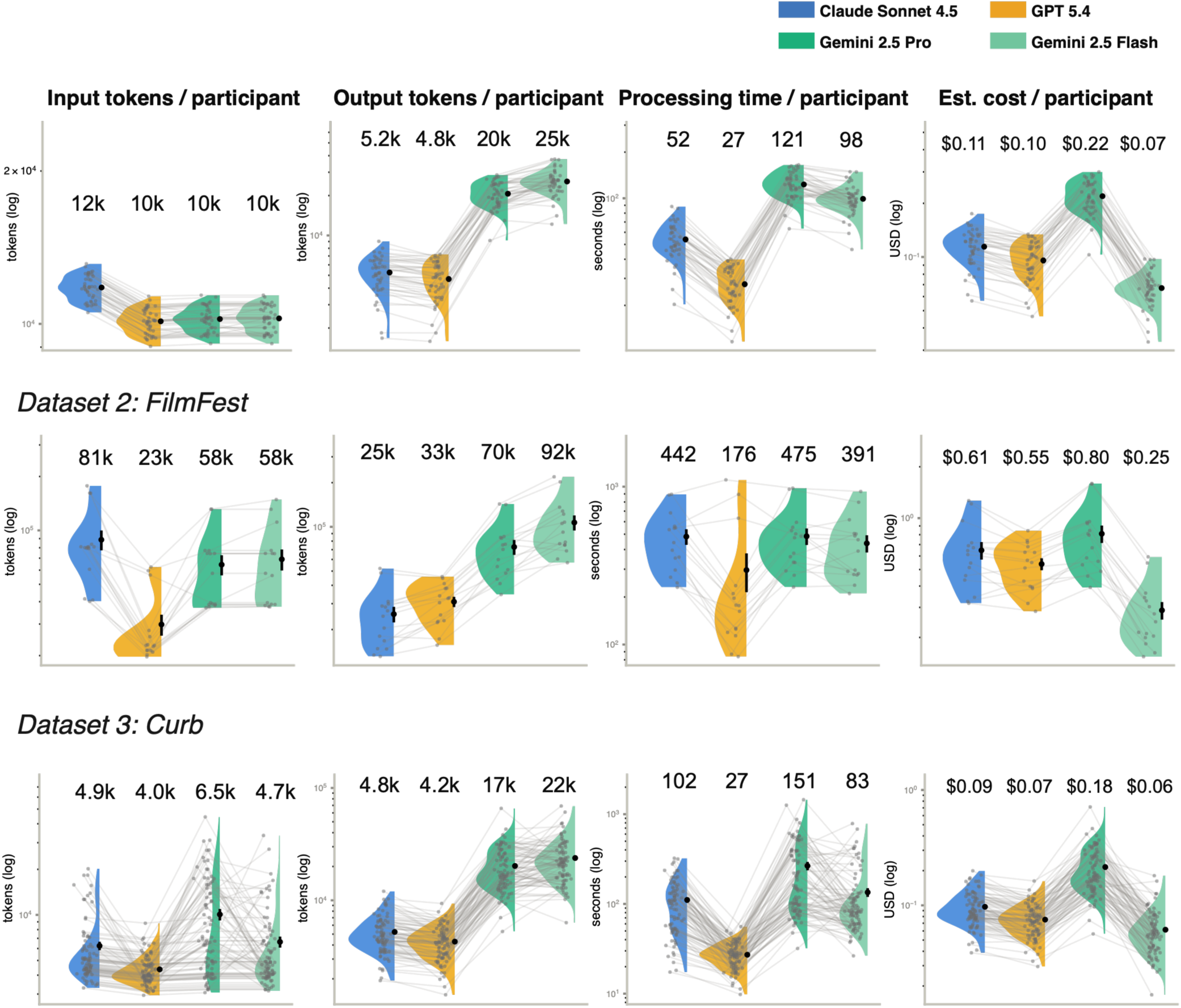
Per-participant model usage in the three datasets. Columns show input tokens, output tokens, processing time, and estimated list-price cost per participant (log scales). Half violins show the distribution across participants, points are individual participants, and faint lines link the same participant across models. Dots and bars show the mean ± SE, and the numbers above each violin are medians. Dataset 1 uses the replication run (n = 43), and Datasets 2 and 3 have 15 and 84 participants. Processing time is missing for one Gemini 2.5 Pro participant in Dataset 3, whose log recorded no per-batch timing.

A failed call is a model response whose JSON output could not be parsed. The pipeline discards such a response and retries the batch in smaller parts. None of the four models had a failed call in the Dataset 1 replication run, and failures were rare in Dataset 2 (Figure S3). The largest cost in Dataset 2 came from six failed GPT-5.4 calls, which used 0.31 million tokens ($2.96 at list price). Most failures occurred in Dataset 3. There, Gemini 2.5 Pro had 217 failed calls (2.45 million tokens, $18.01), Gemini 2.5 Flash had 96 (1.25 million tokens, $2.43), Claude Sonnet 4.5 had 47 (0.28 million tokens, $2.06), and GPT-5.4 had none. For Gemini 2.5 Pro, the failed calls in Dataset 3 cost about as much as all of its successful calls in that dataset ($18.01 vs. $18.05).

**Figure S3.**
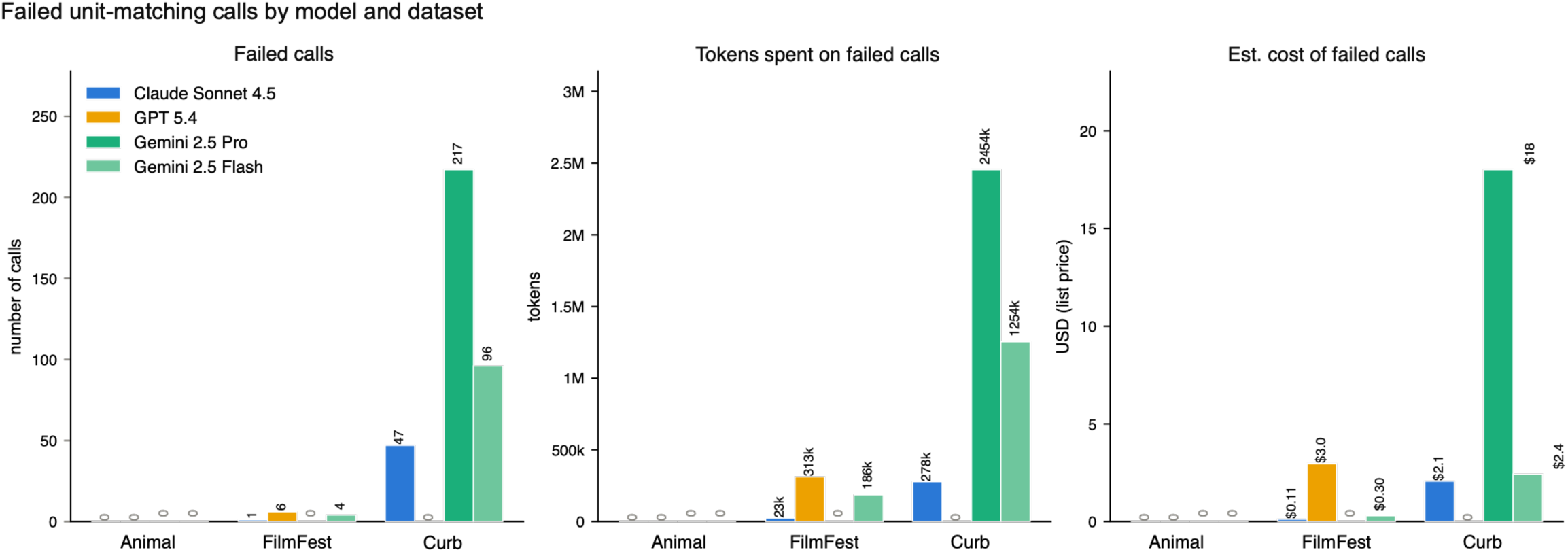
Failed matching calls by model and dataset. Panels show the number of failed calls, the tokens they consumed, and their estimated list-price cost. The Dataset 1 column shows the replication run.

### GPT-5.2

GPT-5.2 (OpenAI) was used in an earlier iteration of the primary comparison and was replaced by GPT-5.4 after OpenAI announced its deprecation (see Method, Models). We report its replication and cost results here and do not report its agreement with human scoring. In the Dataset 1 replication, GPT-5.2 linked 89% of recall units to the same story unit in both runs and assigned the same match category to 86%. Its test–retest agreement (φ = .91, SD = .08; significant for every recall) exceeded the human–human benchmark (t(32) = 9.57, corrected p < .001, BF10 = 1.5 × 10⁸). It was lower than the test–retest agreement of Claude Sonnet 4.5 (t(32) = 10.18, corrected p < .001, BF10 = 6.3 × 10⁸), Gemini 2.5 Pro (t(32) = 3.48, corrected p = .002, BF10 = 23.0), and GPT-5.4 (t(32) = 2.59, corrected p = .019, BF10 = 3.24). It did not differ from the test–retest agreement of Gemini 2.5 Flash (t(32) = 1.73, corrected p = .12, BF10 = 0.71). Test–retest κ was .85 at the information-unit level and .89 at the event-unit level (benchmark, .66 and .77; BF10 = 3.6 × 10¹⁰ and 2.3 × 10⁵). In the replication run, GPT-5.2 had a median cost of $0.08 and a median runtime of 44 s per participant, and it had 14 failed calls.

### Prompts and worked examples

This section reproduces the prompts of the pipeline exactly as they appear in the code (models/autoscorer.py), together with the worked examples of the segmentation prompt (models/proposition_examples.txt). Text in braces is filled in at run time. The {format_instructions} field receives output-format instructions that LangChain generates from the Pydantic schema, {sentences} receives the numbered sentences of a batch, and {example} receives the worked examples. In the matching prompt, {encoding_units} and {recall_units} receive the two unit lists, and {matching_rules} receives the rules for the chosen direction (S3a or S3b). The meta-comment and cross-event rules (S3c–S3e) are included only with the recall as template. The provenance prompt receives, for each match to be resolved, the template unit, the matched unit, and the candidate sentences. The matching and provenance prompts are the versions used in all runs reported here. The segmentation prompt is the version used in the segmentation-consistency analysis, and the Dataset 1 stories and recalls were segmented with an earlier version (Method, Dataset 1).

#### S1. Segmentation prompt (template)

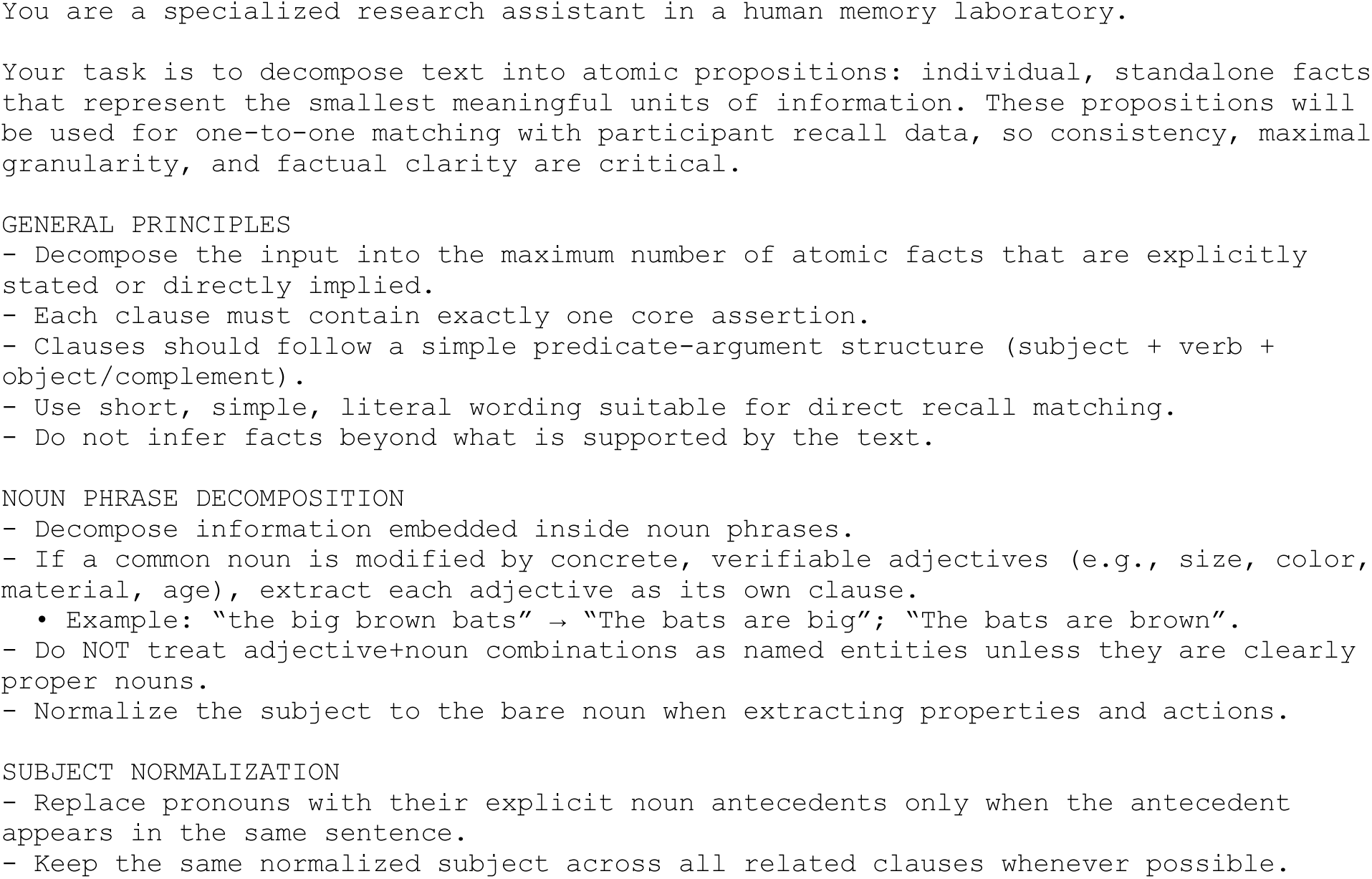
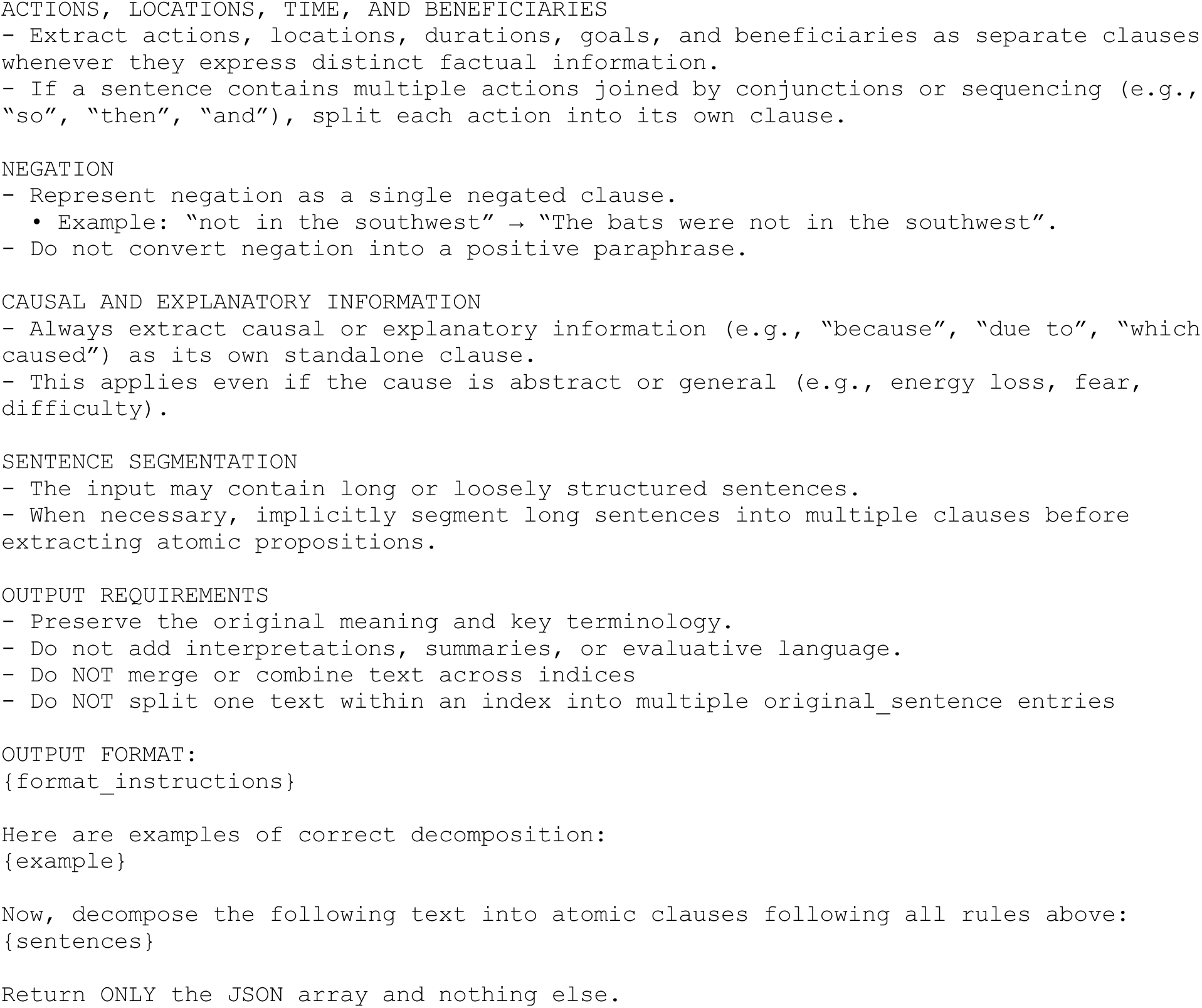

#### S2. Worked examples inserted at {example}

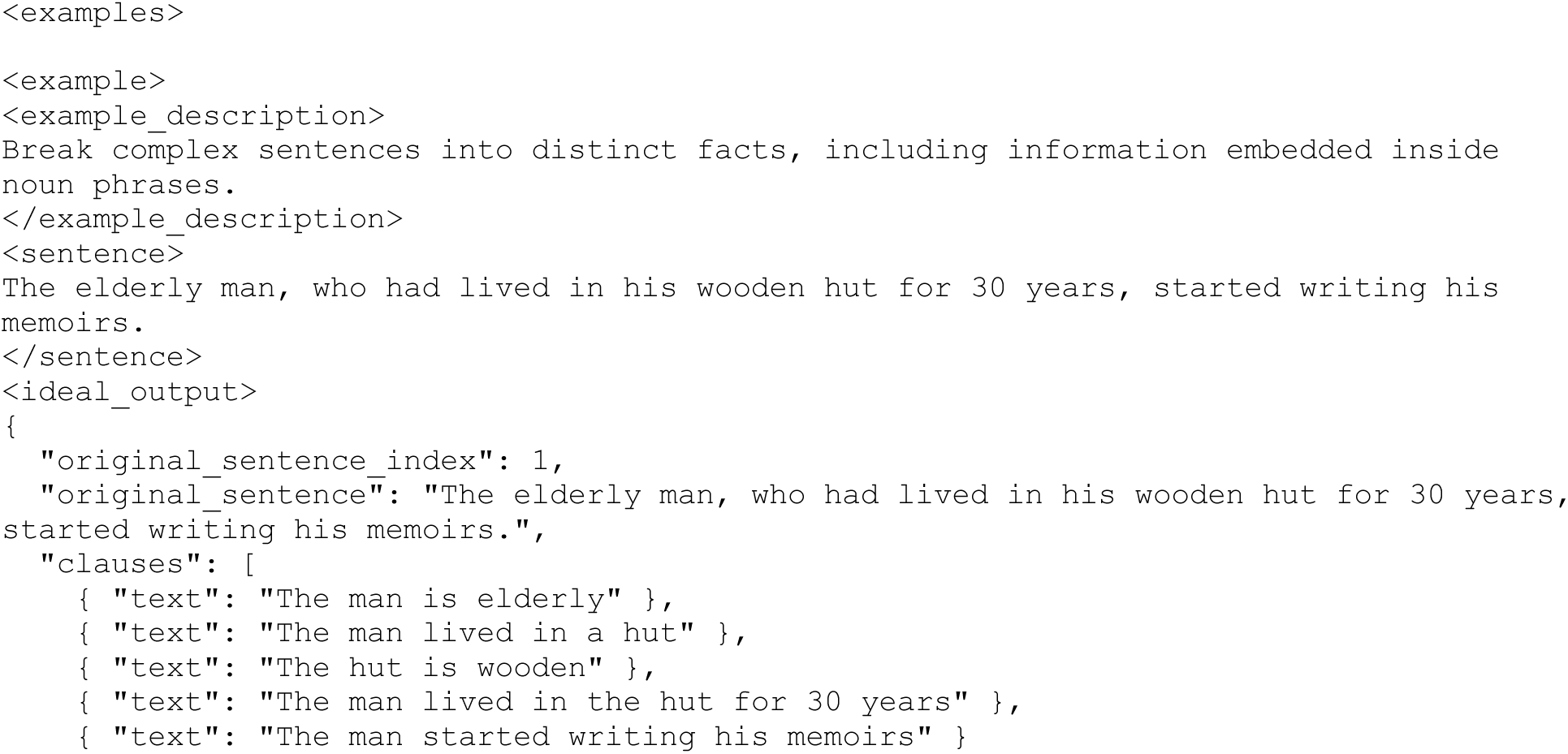
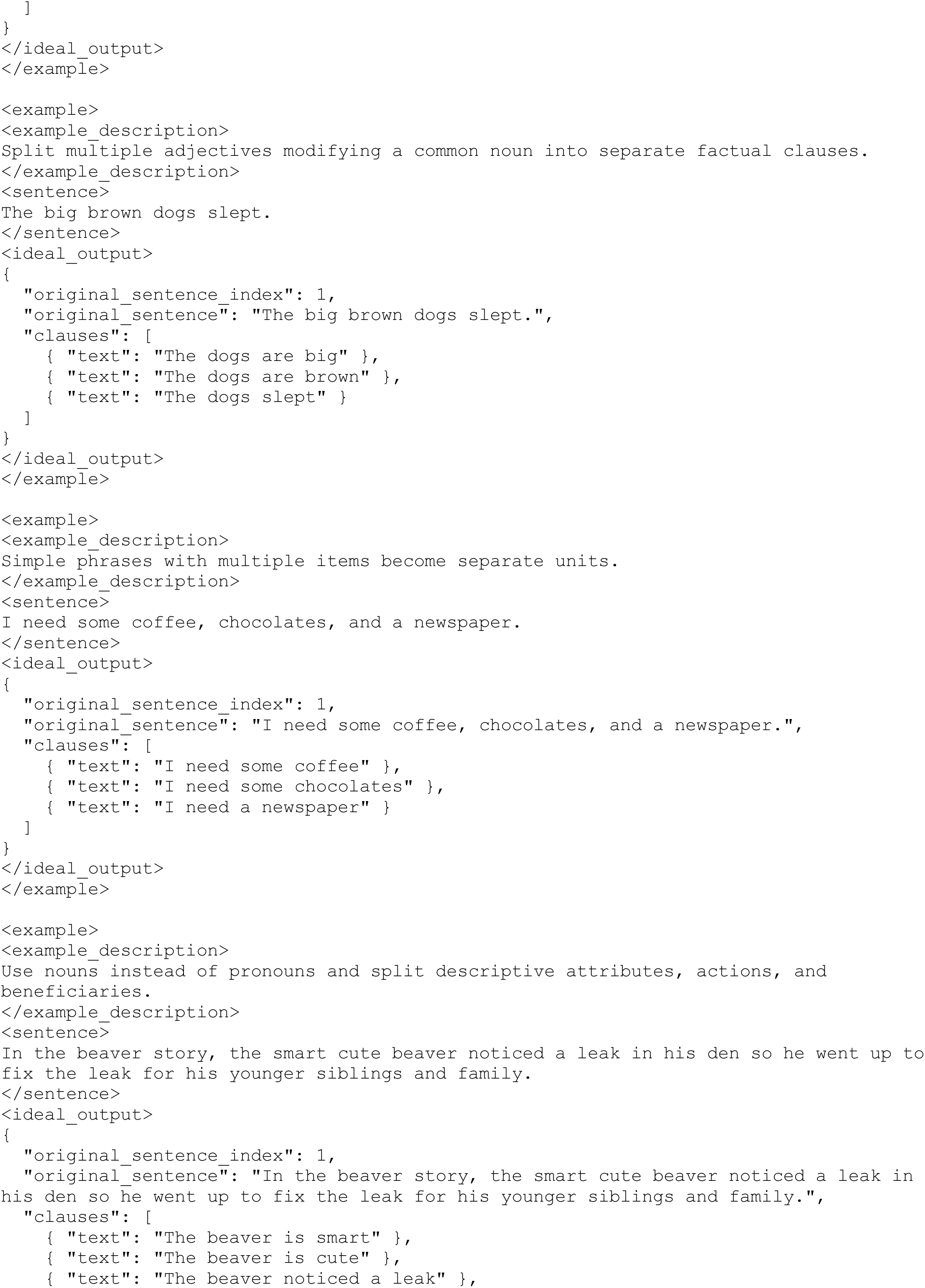
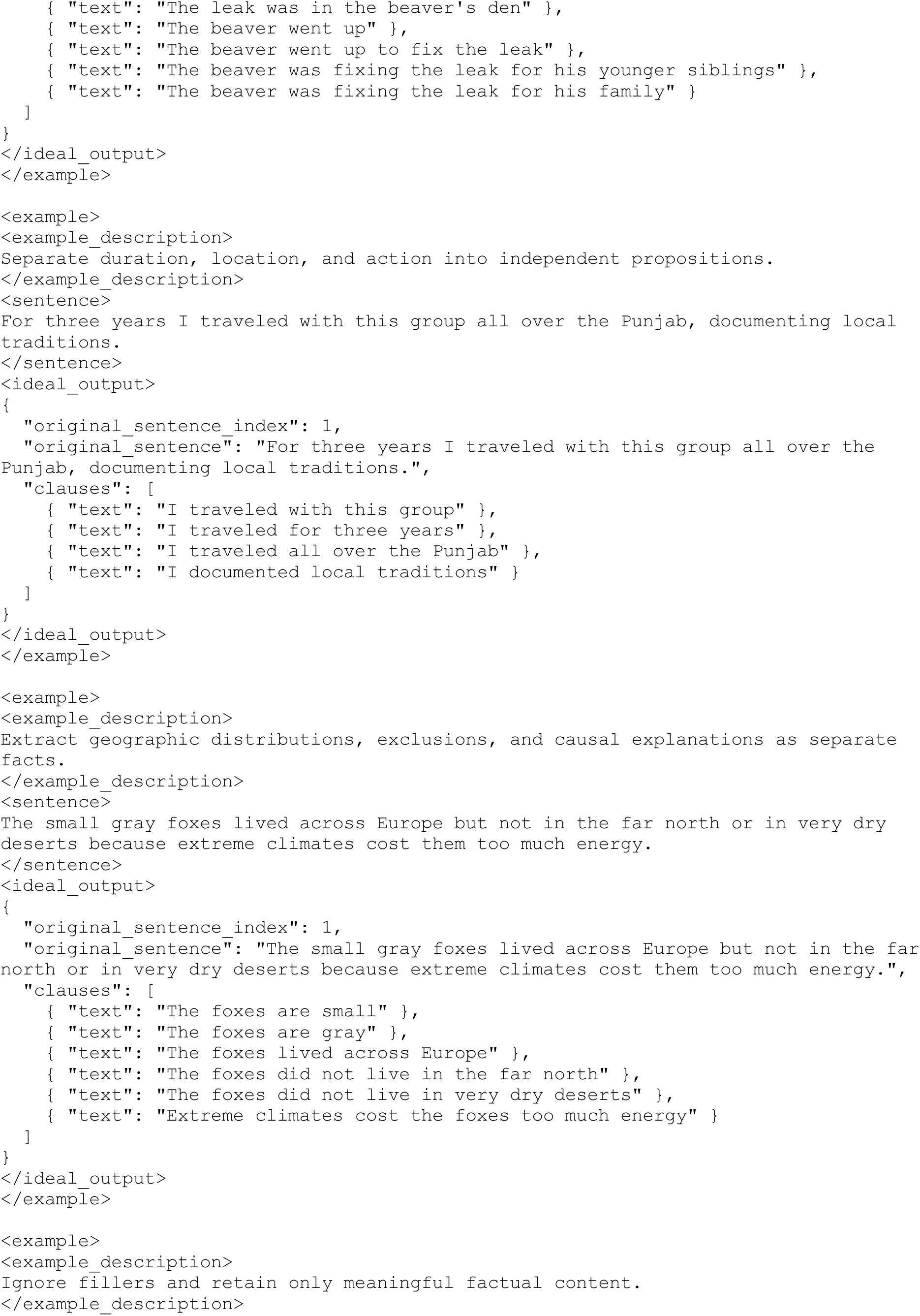
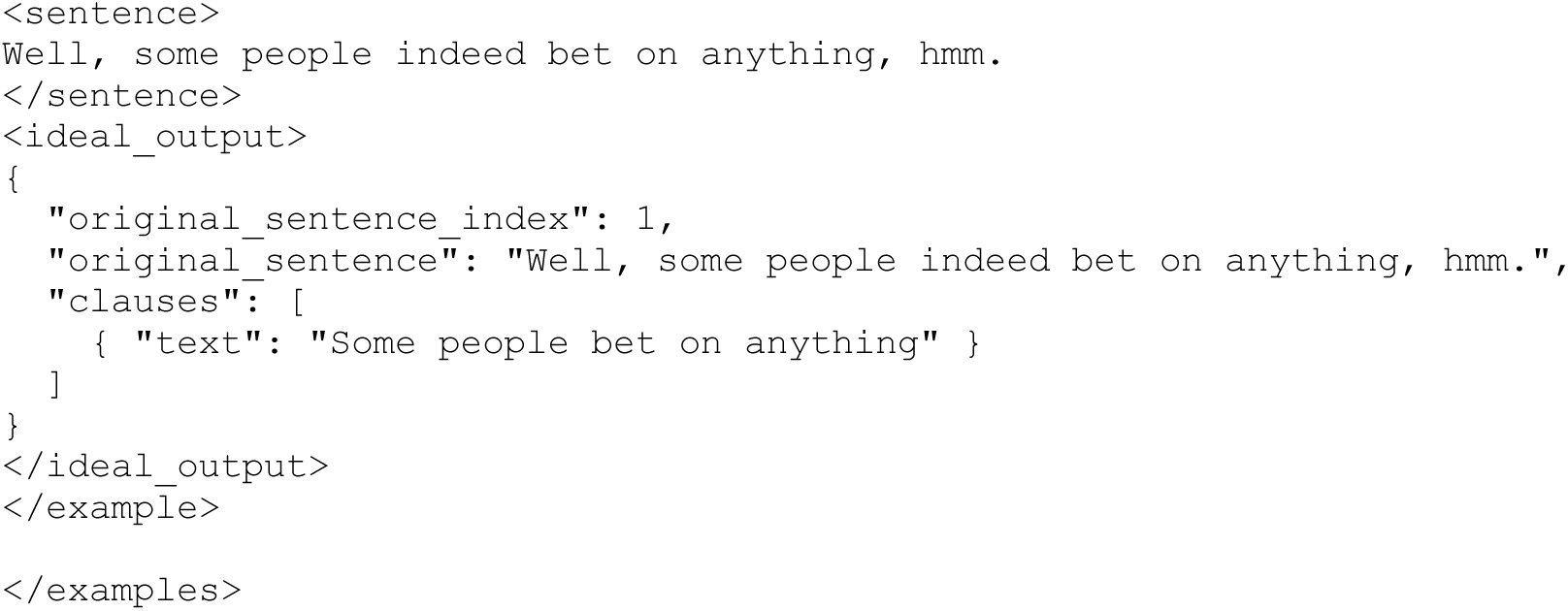

#### S3. Matching prompt (template)

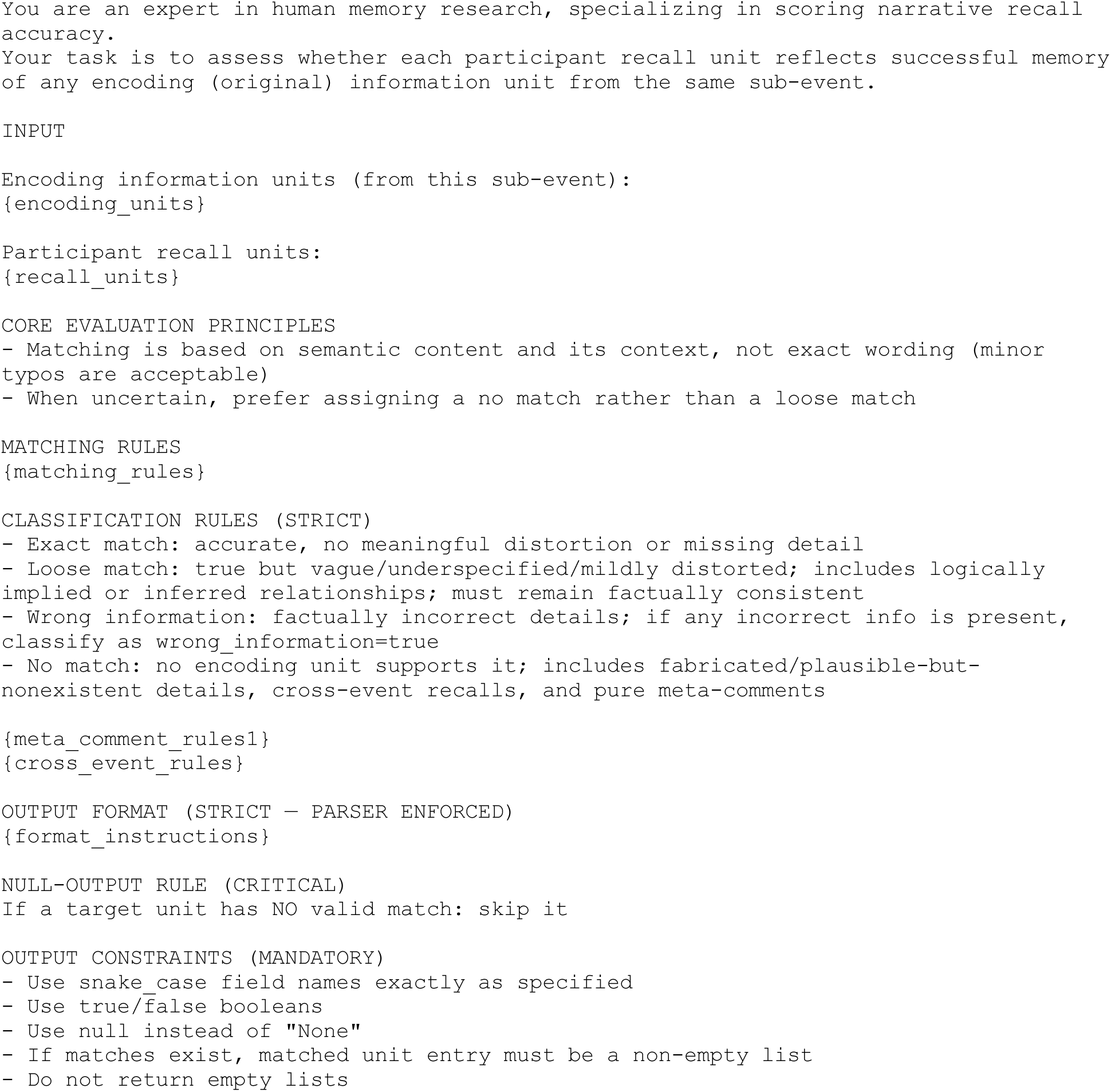
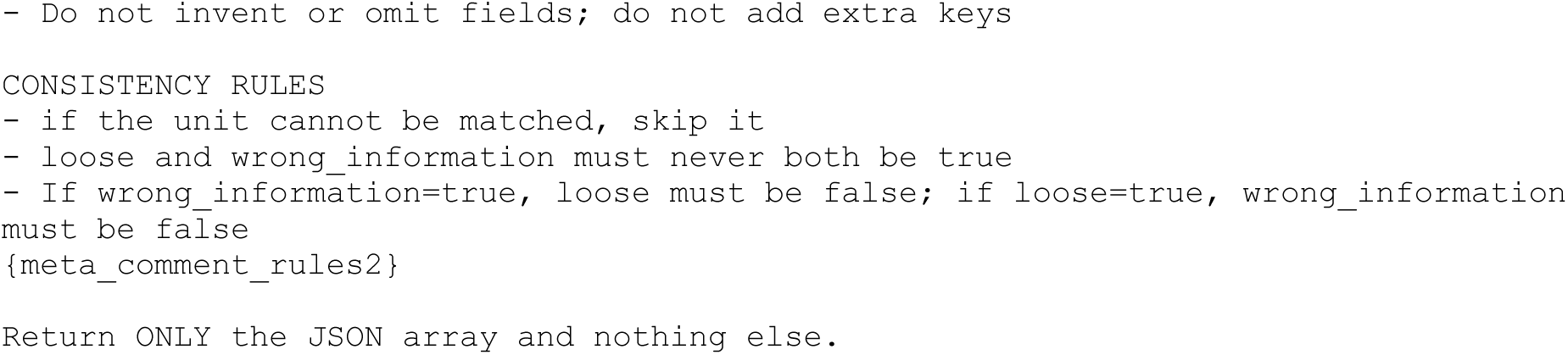

#### S3a. {matching_rules}, recall as template

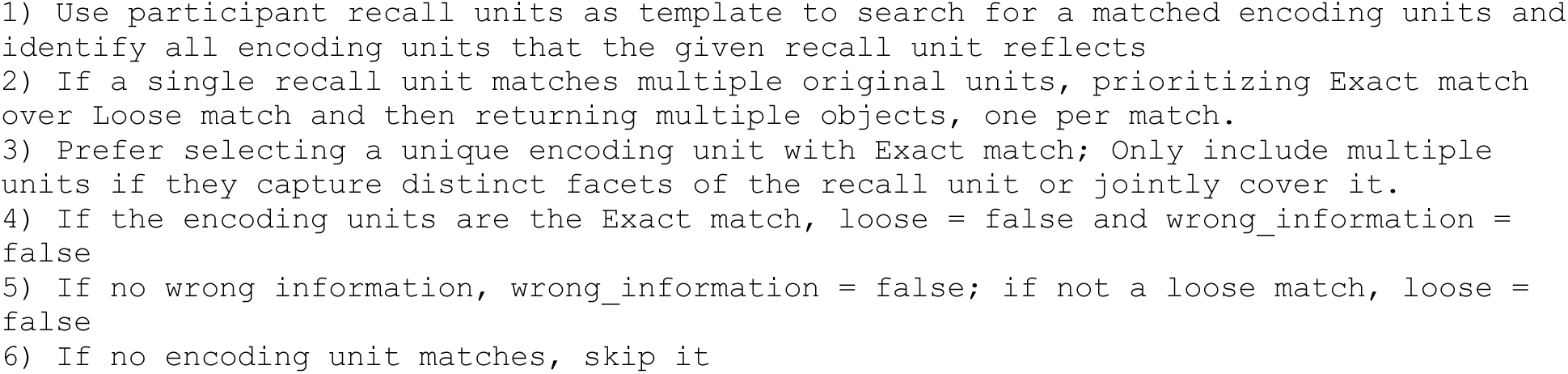

#### S3b. {matching_rules}, story as template

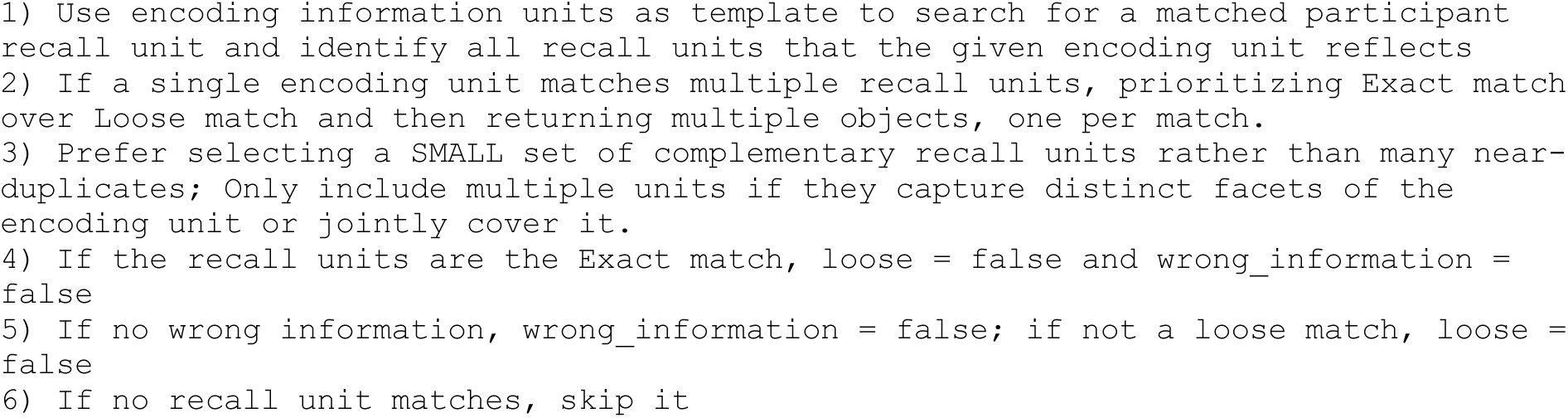

#### S3c. {meta_comment_rules1} (recall as template only)

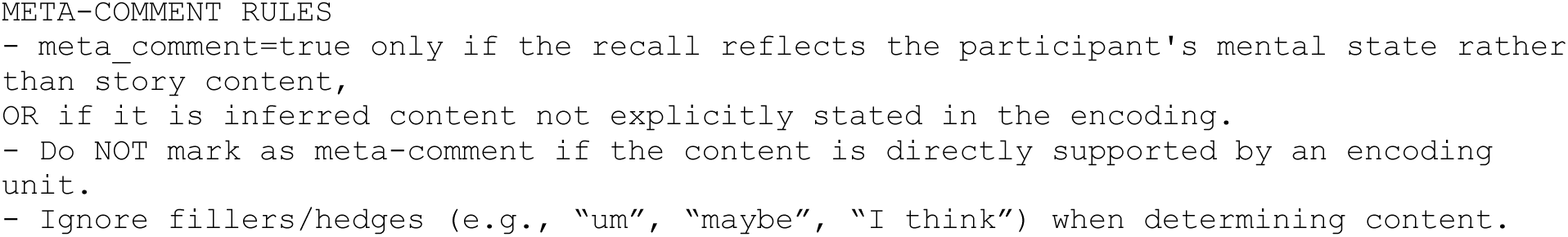

#### S3d. {cross_event_rules} (recall as template only)

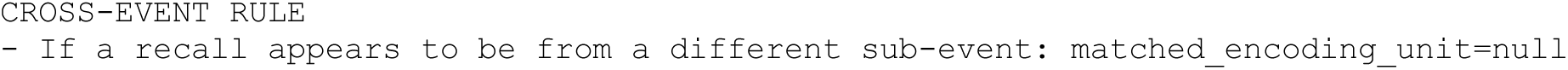

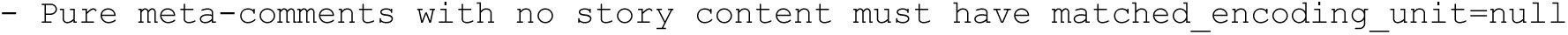

#### S4. Provenance prompt (template)

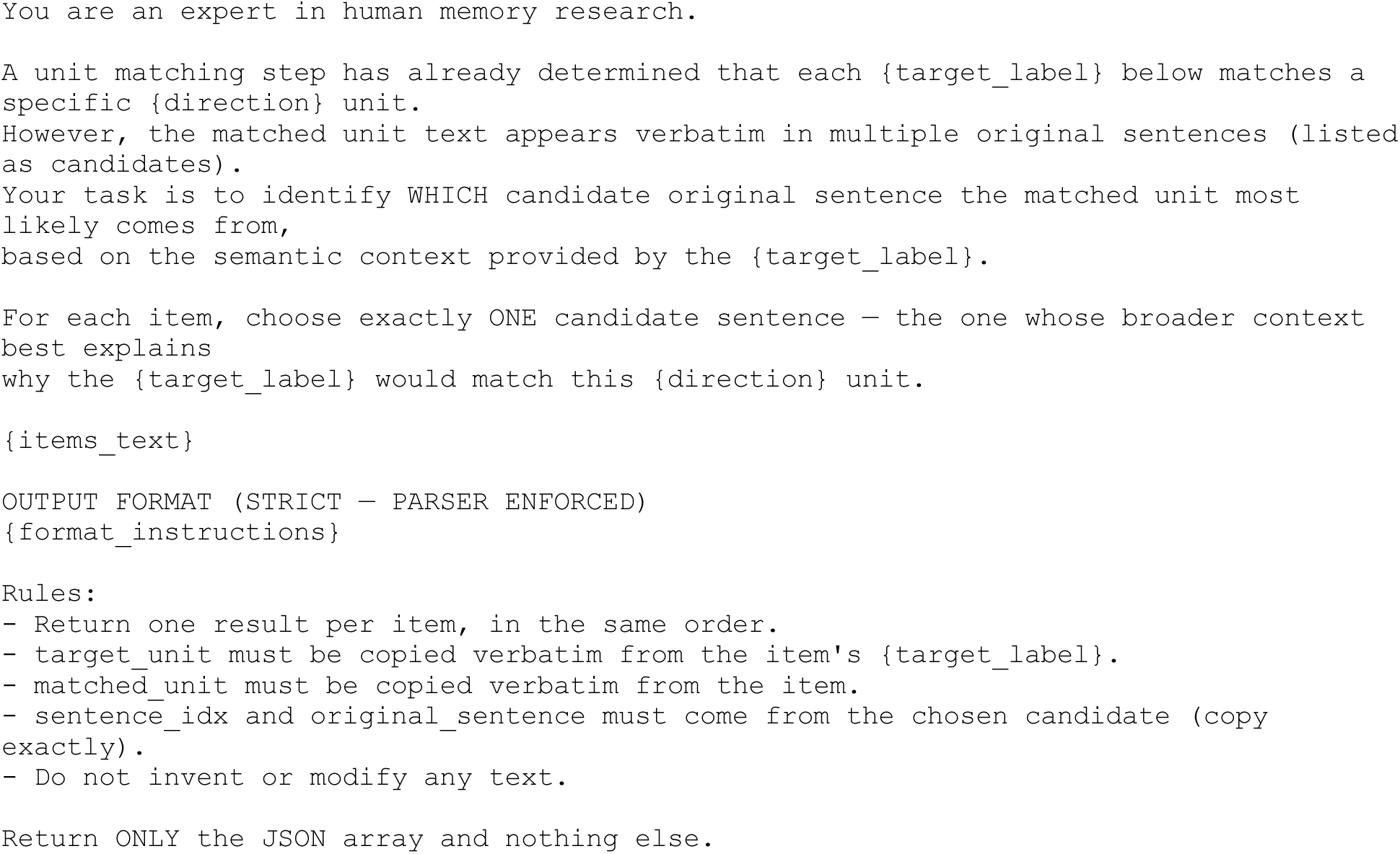

## Notes

### Competing Interest Statement

The authors have declared no competing interest.

